# Modulation of neuronal resilience during aging by Hsp70/Hsp90/STI1 chaperone system

**DOI:** 10.1101/258673

**Authors:** Rachel E. Lackie, Abdul R. Razzaq, Sali M.K. Farhan, Gilli Moshitzky, Flavio H. Beraldo, Marilene H. Lopes, Andrzej Maciejewski, Robert Gros, Jue Fan, Wing-Yiu Choy, David S. Greenberg, Vilma R. Martins, Martin L. Duennwald, Hermona Soreq, Vania F. Prado, Marco A.M. Prado

**Affiliations:** Molecular Medicine, Robarts Research Institute, University of Western Ontario, Canada N6A 5B7; Program in Neuroscience, University of Western Ontario, Canada N6A 5B7; Analytic and Translational Genetics Unit, Center for Genomic Medicine, Massachusetts General Hospital, Harvard Medical School, and The Stanley Center for Psychiatric Research, Broad Institute of MIT and Harvard, Boston, MA, USA 02114; The Edmond and Lily Safra Center for Brain Sciences, Department of Biological Chemistry, The Alexander Silberman Institute of Life Sciences, The Hebrew University of Jerusalem, Jerusalem, Israel 91904; Laboratory of Neurobiology and Stem cells, Department of Cell and Developmental Biology; Institute of Biomedical Sciences, University of Sao Paulo, Sao Paulo Brazil CEP 05508-900; Department of Biochemistry, University of Western Ontario, London, Ontario, Canada N6A 5B7; Department of Physiology and Pharmacology, University of Western Ontario, London, Ontario, Canada N6A 5B7; Department of Medicine, University of Western Ontario, London, Ontario, Canada N6A 5B7; Department of Pathology and Laboratory Medicine, University of Western Ontario, London, Ontario, Canada N6A 5B7; Department of Anatomy & Cell Biology, Schulich School of Medicine and Dentistry, University of Western Ontario, London, Ontario, Canada N6A 5B7; International Research Center, A.C. Camargo Cancer Center, São Paulo, Brazil 01508-010

## Abstract

Chaperone networks are dysregulated with aging and neurodegenerative disease, but whether compromised Hsp70/Hsp90 chaperone function directly contributes to neuronal degeneration is unknown. Stress-inducible phosphoprotein-1 (STI1; STIP1; HOP) is a co-chaperone that simultaneously interacts with Hsp70 and Hsp90, but whose function *in vivo* remains poorly understood. To investigate the requirement of STI1-mediated regulation of the chaperone machinery in aging we combined analysis of a mouse line with a hypomorphic *Stip1* allele, with a neuronal cell line lacking STI1 and in-depth analyses of chaperone genes in human datasets. Loss of STI1 function severely disturbed the Hsp70/Hsp90 machinery *in vivo*, and all client proteins tested and a subset of cochaperones presented decreased levels. Importantly, mice expressing a hypomorphic STI1 allele showed spontaneous age-dependent hippocampal neurodegeneration, with consequent spatial memory deficits. STI1 is a critical node for the chaperone network and it can contribute to age-dependent hippocampal neurodegeneration.

## Introduction

The heat shock proteins 70 (Hsp70) and 90 (Hsp90) are ubiquitously expressed molecular chaperones that promote folding and activation of proteins and are also involved in targeting misfolded or aggregated proteins for refolding or degradation (Lackie et al., 2017). Hsp70 binds indiscriminately to proteins in the early stages of translation and folding and help them to adopt and maintain native conformations. It also prevents aggregation and supports refolding of aggregated and misfolded proteins (Mayer, 2013). Hsp90 is mainly involved in a later stage of activation and supports the maturation and activation of a specific set of “client” proteins, many of which, such as steroid hormone receptors, kinases, and transcription factors, are involved in signaling (Picard, 2006; Taipale et al., 2012; Zhao et al., 2005).

In eukaryotes, both Hsp70 and Hsp90 are regulated by different co-chaperones that tune their activities (Ebong, Beilsten-Edmands, Patel, Morgner, & Robinson, 2016; Harst, Lin, & Obermann, 2005; Hildenbrand et al., 2011; J. Li, Richter, & Buchner, 2011). Client proteins are initially recruited by a complex formed between Hsp70 and its co-chaperone Hsp40 and are then transferred to Hsp90 with the help of the co-chaperone stress-inducible phosphoprotein 1 (See Figure 1, STI1, STIP1 or HOP for Hsp organizing protein in humans). STI1 contains three tetratricopeptide repeat domains (TPR1, TPR2Aand TPR2B); two of them bind to Hsp70 (TPR1 and TPR2B) and one binds to Hsp90 (TPR2A) (Schmid et al., 2012a). STI1 has been shown to physically interact simultaneously with both chaperones and regulate their activity, facilitating the transfer of client proteins (Johnson, Schumacher, Ross, & Toft, 1998; C. T. Lee, Graf, Mayer, Richter, & Mayer, 2012; Rohl, Wengler, et al., 2015; Schmid et al., 2012a).

**Figure 1:**
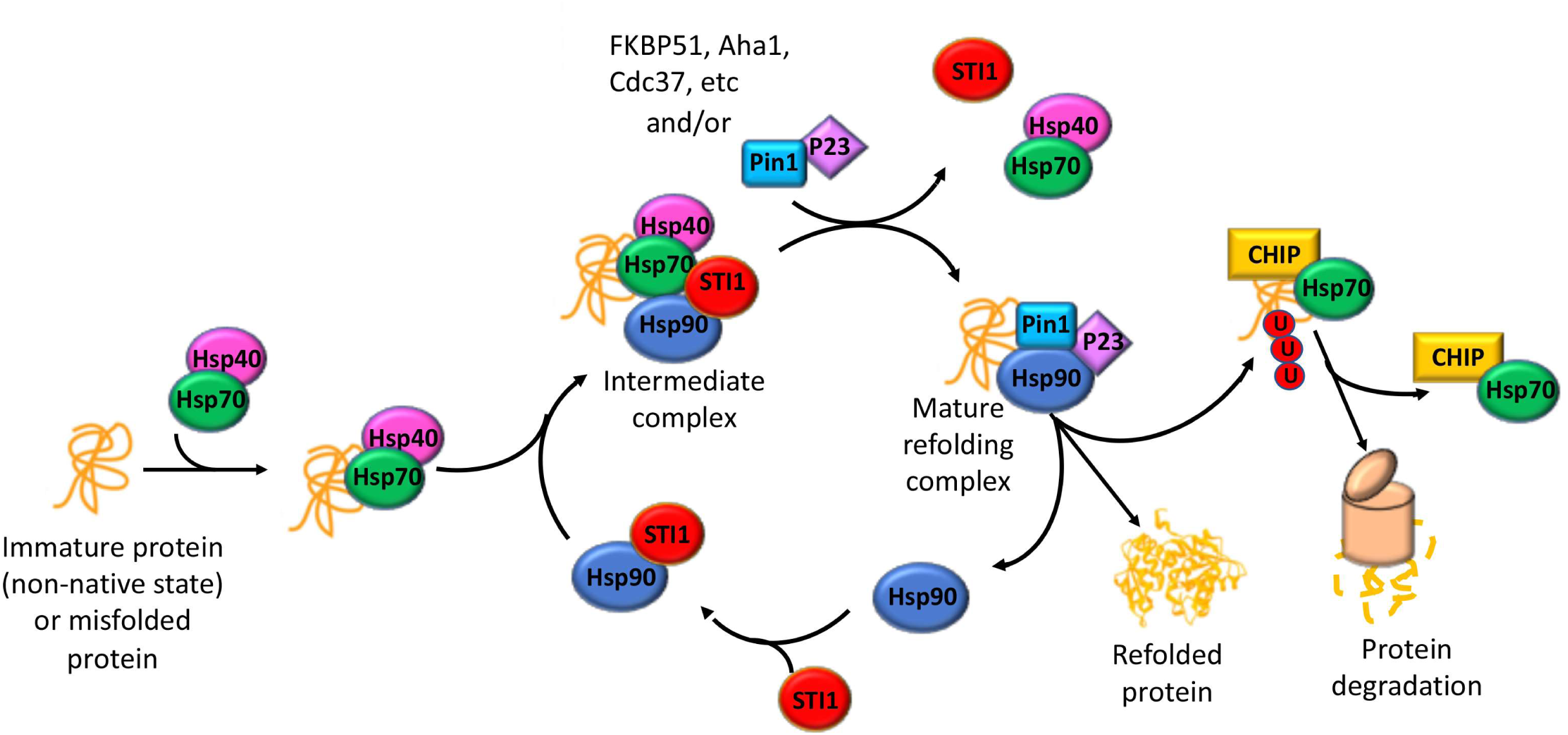
Hsp70/Hsp90 chaperone machinery can support protein maturation or degradation with the help of a number of co-chaperones. This simplified cartoon demonstrates how co-chaperones of Hsp70 and Hsp90 help coordinate the transfer of client proteins to become mature, functional proteins. CHIP and STI1 compete for Hsp70 binding and STI1 helps to mature clients by physically linking Hsp70 and Hsp90, allowing client transfer from Hsp70 to Hsp90. Depending on client, or if the client is being marked for degradation, a variety of other co-chaperones other than STI1 can be employed by this machinery.

Besides its role in the client folding pathway, STI1 has been suggested to be involved in a number of different functions including shuttling of some proteins from the cytosol to the mitochondria (Hoseini et al., 2016); facilitating gene transcription by removing nucleosomes at target promoters (Floer, Bryant, & Ptashne, 2008); and maintaining genome integrity by silencing transposons (Gangaraju et al., 2011; Karam, Parikh, Nayak, Rosenkranz, & Gangaraju, 2017). Furthermore, STI1 can be secreted by different cells, including astrocytes and microglia, and binding of extracellular STI1 to the prion protein triggers pro-survival signaling cascades and prevents Aβ toxicity in neurons (Linden et al., 2008; Lopes et al., 2005; Ostapchenko et al., 2013; Zanata et al., 2002).

Noteworthy, yeast cells null for STI1 are viable under optimal conditions (Chang, Nathan, & Lindquist, 1997; Y. Song & Masison, 2005), but they are highly sensitive to Hsp90-inhibiting compounds and grow poorly under limiting conditions (Chang et al., 1997; Y. Song & Masison, 2005). In worms STI1 lacks the TPR1 domain, but it is still functional and connects Hsp70/Hsp90 (Y. Song & Masison, 2005). *C. elegans* lacking STI1 are viable but are less resilient to stress and display reduced lifespan (Gaiser, Brandt, & Richter, 2009b). In contrast, knockout of STI1 in mice leads to embryonic lethality (Beraldo et al., 2013), indicating that in mammals the roles of STI1 are essential for life and cannot be compensated by other proteins. To note, during mouse development, STI1 co-chaperone activity seems to be a critical mechanism for the modulation of apoptosis and cellular resilience (Beraldo et al., 2013).

The chaperone machinery is essential for protein quality control and is thought to be particularly important in neurodegenerative disorders such as Alzheimer’s, Parkinson’s and Huntington’s disease (Fontaine et al., 2016; Pratt, Gestwicki, Osawa, & Lieberman, 2015). In yeast, STI1 interaction with Hsp70 can redirect toxic amyloid-like proteins into cytosolic foci, thus increasing cellular viability (Wolfe, Ren, Trepte, & Cyr, 2013). Remarkably, recent experiments in yeast suggest that excessive demand for chaperone activity can disturb chaperone function by decreasing STI1 interaction with members of the chaperone network (Farkas et al., 2018). Dysregulation of the Hsp70/Hsp90 chaperone network transcriptome is present in aging brains and in neurodegenerative disease (Brehme et al., 2014). However, whether STI1 and Hsp70/Hsp90 are required *in vivo* to maintain homeostasis in mammals during aging is unknown. To address this critical question, we engineered a mouse line with a STI1 hypomorphic allele that retained partial functionality allowing survival of mice to adulthood. We combined analyses of human genetic datasets, this novel hypomorphic mouse line and a CRISPR-Cas9 STI1 knockout neuronal cell line to understand the requirement of STI1 for the functionality of chaperone networks. Our experiments reveal that limiting STI1 function due to the hypomorphic STI1 allele strongly reduces neuronal resilience during aging, suggesting a mechanism by which compromised chaperone network function may contribute to neurodegeneration.

## Results

### The ΔTPR1 hypomorphic allele is expressed at low levels but is sufficient for mouse survival

To investigate the relationship between STI1 and the chaperone network in mammals, we generated hypomorphic TPR1-deprived STI1 mice using the Cre/lox system to remove exons 2 and 3 of the STI1 gene (Fig. 2A). We focused on removing the TPR1 domain because STI1 TPR domains are well conserved from yeast to humans and the TPR1 domain is absent in *C. elegans*, yet the protein can still regulate Hsp70/Hsp90 (Gaiser et al., 2009b). Moreover, we confirmed the recombination of the *Stip1* locus by genome sequencing (data not shown, see STI1-flox and ΔTPR allele cartoon in Figure 2A). After recombination, we confirmed that the alternative translation initiation codon of the mutated mRNA lacks a neighboring Kozak consensus (Kozak, 1986), which likely contributed to less efficient translation of ΔTPR1 protein.

**Figure 2:**
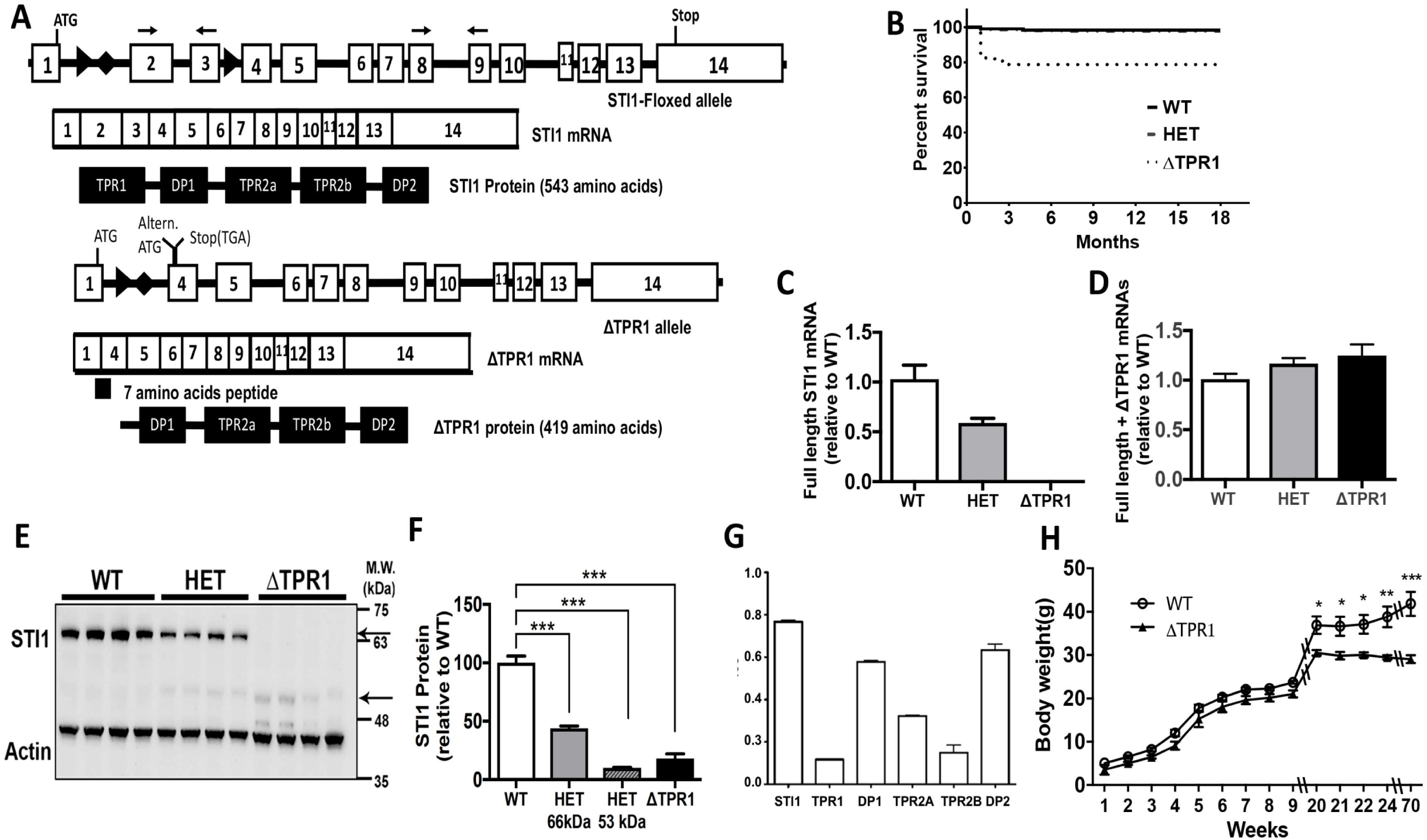
Generation and characterization of ΔTPR1 mice. **A.** Simplified cartoon of the *Stip1* gene locus with or without recombination to remove exons 2 and 3 and the major domains of the STI1 protein with or without the TPR1 domain (143 amino acid deletion). **B.** Percent survival of male WT, ΔTPR1 heterozygous and ΔTPR1 homozygous mice from birth to final age of collection, 18 months old. (Solid black line, WT; HET smaller black dash with dots; ΔTPR1 small black dots). **C.** Analysis of adult (15-18 months old) cortical brain tissue STI1 transcripts using primers for exons 2 and 3 (see primer location on A), which amplify full length STI1 mRNA but not ΔTPR1 mRNA. **D.** qPCR analysis for STI1 transcripts using primers for exons 8 and 9 (see primer location on A, N = 8) which amplify both full length and ΔTPR1 mRNA. **E.** Representative Western blots of STI1 expression in adult cortical brain extracts. Arrow full-length STI1, arrowhead ΔTPR1 protein. **F.** Quantitative analysis of STI1 levels in WT (white bar), ΔTPR1 heterozygous (WT/ΔTPR1: HET) for full length STI1 (66 kDa, light grey bar) and truncated protein (53 kDa, dark grey bar with stripes) and homozygous ΔTPR1 mice (black bar). Data are mean ± SEM (N = 8). One-way ANOVA with corrections for multiple comparisons with Dunnett’s test. **G.** Surface plasmon resonance analysis of affinity of STI1 antibody for each domain of STI1. **H.** Comparison of body weights of mice from 1-70 weeks of age (closed circle for WT and dark triangle for ΔTPR1 mice). N = 7-10 mice for each time point. Data analyzed using Two-Way ANOVA, multiple comparisons corrected with Sidak’s test. WT (WT/WT), HET (WT/ΔTPR1), ΔTPR1 (ΔTPR1/ΔTPR1). Data are mean ± SEM (*p < 0.05, **p < 0.001, ***p < 0.0001). **J.**

Homozygous mutant STI1 mice (ΔTPR1 mice) were viable but they were born on a significant lower frequency than expected from a Mendelian distribution. That is, out of 488 pups born from breeding WT/ΔTPR1 to WT/ΔTPR1 mice, 154 (32%) were WT, 277 (57%) were WT/ΔTPR1, and 57 (12%) were ΔTPR instead of 25%, 50% and 25% respectively (χ2 = 47.49, df = 2, p < 0.0001). Interestingly, at embryonic day E17.5, the proportion of homozygous ΔTPR1 mutants was close to the expected Mendelian frequency of 25% (out of 33 pups 3 (9%) were WT, 22 (67%) were WT/ΔTPR1, and 8 (24%) were ΔTPR1; χ2 = 5.182, df = 2, p = 0.0075), suggesting that the decreased Mendelian distribution we observed for ΔTPR1 mice was because pups were dying immediately after birth. Supporting this hypothesis, ΔTPR mice showed a significantly higher mortality rate during the first month and 17.5% of the cohort died before 30 days of age, compared to 0.5% of WT/ΔTPR1 and 1.0% of the WT siblings. Interestingly, survival rate of ΔTPR1 mice that lived through and after the first month was not different from that of WT/ΔTPR1 and WT mice (Fig. 2B).

Strikingly, one *ΔTPR1* allele was able to rescue the early embryonic lethality of STI1 null mutants. Specifically, we bred ΔTPR1 females to heterozygous null mice for STI1 (STI1 WT/KO mice, (Beraldo et al., 2013) and observed that out of 21 pups born alive, two were ΔTPR1/STI1KO (9.5%) while 19 were ΔTPR1/WT (90.5%) instead of 50% and 50% as expected from the Mendelian distribution. One of the ΔTPR1/STI1KO pups died one day after birth and the other survived to adulthood. These data suggest that STI1 lacking TPR1 has the necessary ability to allow mammalian development to proceed. Because of their frailty we did not further proceed to obtain mice with only one ΔTPR1 allele.

We used qPCR to determine mRNA expression for the mutated locus (Fig. 2C and D). Analysis with primers targeting exons 2 and 3 (Fig. 2A, these primers detect full length STI1 mRNA, but not ΔTPR1-STI1 mRNA) showed 50% reduction in full-length STI1 mRNA levels in heterozygous mutants, whereas in homozygous mutants, full length STI1 mRNA was not detected in adult cortical tissue (Fig. 2C). Importantly, primers flanking exons 8 and 9 (Fig. 2A, which detect both full length and ΔTPR1 STI1 mRNA) revealed that expression level of ΔTPR1 STI1 mRNA in ΔTPR1 mice was like that of STI1 full length mRNA in WT mice (Fig. 2D, one-way ANOVA, p = 0.1022).

Immunoblot analysis demonstrated that ΔTPR1 mice lacked full length STI1 protein (66 kDa) and instead expressed a truncated protein with reduced molecular mass of 53 kDa (Fig. 2E). The deleted TPR1 domain is predicted to be 12-13 kDa in size, hence the apparent molecular mass of the mutant protein equals the predicted molecular mass of STI1 (66 kDa) minus 13 kDa. We detected close to 80% reduction in mutant protein levels in ΔTPR1 mice when compared to WT littermates (Fig. 2F, p < 0.0001). Control experiments demonstrated that our polyclonal antibody recognizes several epitopes on the STI1 protein, therefore excluding the possibility that the reduced levels of immunostaining observed were due to decreased binding of the antibody to deleted epitopes (Figure 2G).

Two possibilities could explain decreased STI1 levels in ΔTPR1 mice: 1) the ΔTPR1 protein is unstable and therefore undergoes rapid degradation or 2) the ΔTPR1 mRNA is poorly translated. Because the yeast ΔTPR1 protein is stable (Rohl, Tippel, et al., 2015; Rohl, Wengler, et al., 2015; Schmid et al., 2012b) and the *C. elegans* protein naturally lacks the TPR1 domain (Gaiser, Brandt, & Richter, 2009a; H. O. Song et al., 2009), it is unlikely that decreased levels of the ΔTPR1 protein in mice is a consequence of instability. On the other hand, the mRNA generated after deletion of exons 2 and 3 was expected to be poorly translated. The 5’ end of the STI1 mRNA, including the translational initiation codon, was preserved in the ΔTPR1 mRNA, but a UGA stop codon was created 18 nucleotides downstream of the initiation codon. Thus, only a 7 amino acid peptide would be expected to be generated when this canonical initiation codon is used. However, overlapping the newly created UGA codon there is an AUG that can work as an alternative initiation codon and generate the ΔTPR1 protein (Fig. 2A). Much less ΔTPR1 protein is expected to be generated because of low efficacy of the alternative initiation codon. Sequencing analysis of the mutated *Stip1* locus and mRNA confirmed these changes. Regardless of the mechanism, the hypomorphic STI1 mice have a truncated STI1 protein that is expressed at low levels but provides sufficient activity for survival.

Noteworthy, while young ΔTPR1 mice and WT littermate controls showed similar weight, adult ΔTPR1 mice gained less weight than WT littermate controls (Fig. 2H). To determine whether ΔTPR1 mice showed any metabolic phenotype, we tested them on metabolic cages at 15-18 months of age. Mice were habituated to the metabolic cages for 16 h and data were collected over the following 24 h (Table 1). We observed that food intake was not significantly different between ΔTPR1 mice and littermate controls, both during the light and the dark phases of the day, indicating that difference in weight was not due to decreased food intake. Likewise, water consumption was similar between both genotypes. On the other hand, ΔTPR1 mice showed increased ambulatory daily activity when compared to WT littermate controls, mainly due to increased locomotion during the light cycle. Also, ΔTPR1 mice showed a significant increase in total activity, which includes not only locomotion but also grooming, sniffing, tail flicking and rearing, both during the light and dark cycle (Table 1). The increased physical activity could explain the decreased body weight observed in ΔTPR1 mice. Also, it could explain the higher volumes of oxygen consumption and carbon dioxide release observed in ΔTPR1 mice during both the light and dark cycles (Table 1). Importantly, the elevated metabolic rate was not a consequence of increased production of heat [Energy expenditure (EE); Table 1]. Furthermore, no significant differences in respiratory exchange rate (RER), a parameter that reflects the relative contributions of carbohydrate and fat oxidation to total energy expenditure, were observed between ΔTPR1 mice and littermate controls.

**Table 1.** Metabolic parameters of mutant mice and littermate controls

|  | Light |  | Dark |  |
| --- | --- | --- | --- | --- |
|  | WT (N=) | ΔTPR1 (N=8) | WT (N=7) | ΔTPR1 (N=8) |
| Food intake (g) | 1.137 ± 0.24 | 1.42 ± 0.22 | 1.64 ± 0.25 | 2.10 ± 0.29 |
| Water (ml) | 1.15 ± 0.24 | 1.34 ± 0.22 | 2.15 ± 0.24 | 2.45 ± 0.16 |
| Amb. act. (counts/h) | 366.07 ± 56.47 | 744.93 ± 149.93* | 2197.63 ± 210.08 | 4666.06 ± 1139.59 |
| Total act. (counts/h) | 1201.36 ± 146.92 | 1809.30 ± 257.74* | 4445.30 ± 413.56 | 7904.40 ± 1617.05 |
| VO2 (mL/kg/h) | 1959.53 ± 75.42 | 2544.97 ± 979.28 | 2284.56 ± 47.09 | 3183.77 ± 95.91 |
| VCO2 (mL/kg/h) | 1679.03 ± 113.34 | 2321.37 ± 122.71 | 2047.50 ± 71.89 | 3005.79 ± 90.35 |
| EE (kcal/h) | 0.42 ± 0.02 | 0.38 ± 0.02 | 0.50 ± 0.02 | 0.48 ± 0.01 |
| RER | 0.85 ± 0.03 | 0.91 ± 0.03 | 0.90 ± 0.02 | 0.95 ± 0.03 |
| Body weight | 44.24 ± 1.47 | 30.54 ± 0.92**** |  |  |
| Sleep time | 432.14 ± 16.74 | 400.63 ± 21.71 | 250.00 ± 16.22 | 165.25 ± 23.81 |

Ambulatory activity (Amb. act.) and Total activity (Total act.). EE, energy expenditure; RER, respiratory exchange ratio.
\*p < 0.05 and \*\*\*\*p<0.0001 (WT vs. ΔTPR1).

### ΔTPR1-STI1 does not affect mRNA and protein levels of Hsp70 and Hsp90 but affects a subset of Hsp70-Hsp90 interacting proteins and co-chaperones

We tested whether Hsp90 and Hsp70 mRNA levels were changed in ΔTPR1 mice but did not observe any difference among genotypes (Fig. 3A-B; Hsp90, p = 0.18; and for Hsp70, p = 0.37). We then tested whether ΔTPR1-STI1 affects expression levels of Hsp90 and Hsp70 proteins in aged (15-18-month-old) mice (Fig. 3C-F). We did not find any changes in Hsp90 protein levels (Fig. 3C-E, p = 0.076 for pan Hsp90; p = 0.465 for Hsp90β). There was also no significant difference in Hsp70 levels when we compared controls and ΔTPR1 brain tissues (Fig. 3C and F, p = 0.076). Immunofluorescent labelling for STI1, Hsp70 and Hsp90 revealed that localization of Hsp90 (Fig. 3G) and Hsp70 (Fig. 3H) in ΔTPR1 MEFs (passage 4) was not altered when compared to wild-type MEFs and there was no significant change in STI1 co-localization with these chaperones. Additionally, immunofluorescence confirmed that ΔTPR1 levels in MEFs were reduced.

**Figure 3.**
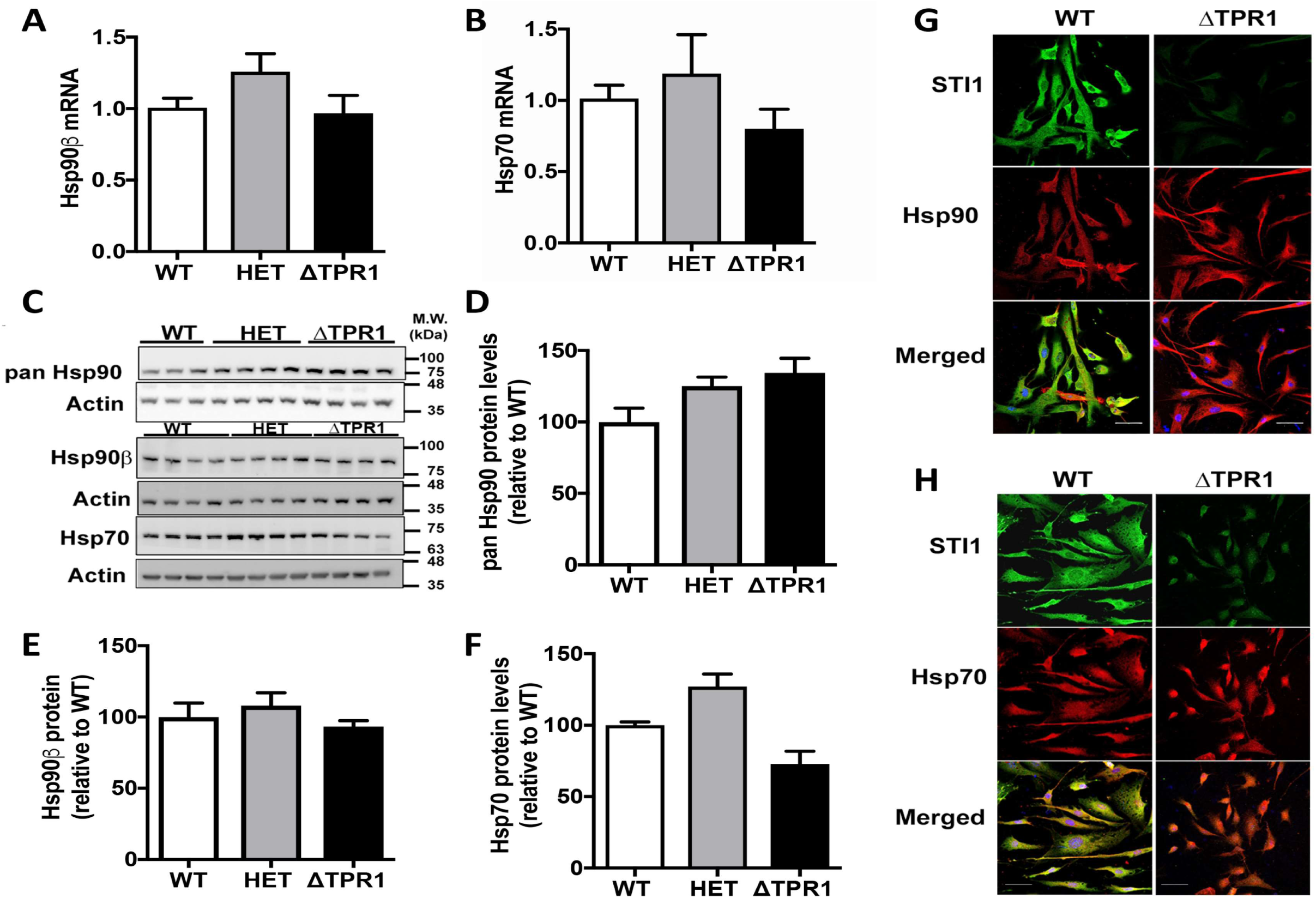
Hsp70 and Hsp90 is unaltered in ΔTPR1 brain tissue and MEFs. **A.** Hsp90β mRNA expression in adult cortical brain tissue. **B.** Hsp70 mRNA expression in adult cortical brain tissue. (N = 4). **C.** Representative Western Blots for Hsp90 and Hsp70 expression in male adult mice cortical tissue (N = 4/genotype). **D-F.** Quantification of total Hsp90, Hsp90β or Hsp70 protein levels in WT, HET and ΔTPR1 cortical tissue. **G.** Immunofluorescence analysis of the localization of Hsp90 (red); **H.** Hsp70 (red) and STI1 (green) in MEFs. Scale bar = 50 µm.

We also investigated the levels of different regulators of the heat shock response and co-chaperones known to interact with Hsp70 and Hsp90 in aged mice (Fig. 4). Levels of the transcription factor HSF1, a major regulator of the heat shock response that is modulated by Hsp90 (Dai et al., 2003), were not affected in ΔTPR1 brain tissue (Fig. 4A, B, p = 0.30). Likewise, levels of Hsp40 (Fig. 4A, C), a DnaJ protein that is a co-chaperone for Hsp70 and is present in the Hsp70-STI1 complex (Cyr, Lu, & Douglas, 1992; Frydman, Nimmesgern, Ohtsuka, & Hartl, 1994; Morgner et al., 2015; Tsai & Douglas, 1996) was not significantly changed (p = 0.61). Levels of the peptidyl-prolyl isomerase (PPIase) FKBP51, an Hsp90 co-chaperone that is involved in the maturation of steroid hormone receptors and stabilization of Tau species (Barent et al., 1998; Dickey et al., 2007; Jinwal et al., 2010; Nair et al., 1997) was also normal (Fig. 4D and E, p = 0.548). Additionally, levels of the co-chaperones Aha1 (p = 0.28), p23 (p = 0.311) and Cdc37 (p = 0.15) were not affected in STI1 hypomorphic mouse tissue (Fig. 4F-I). These results indicate that important players in the Hsp70/Hsp90 chaperone network were not affected in ΔTPR1 mice.

**Figure 4.**
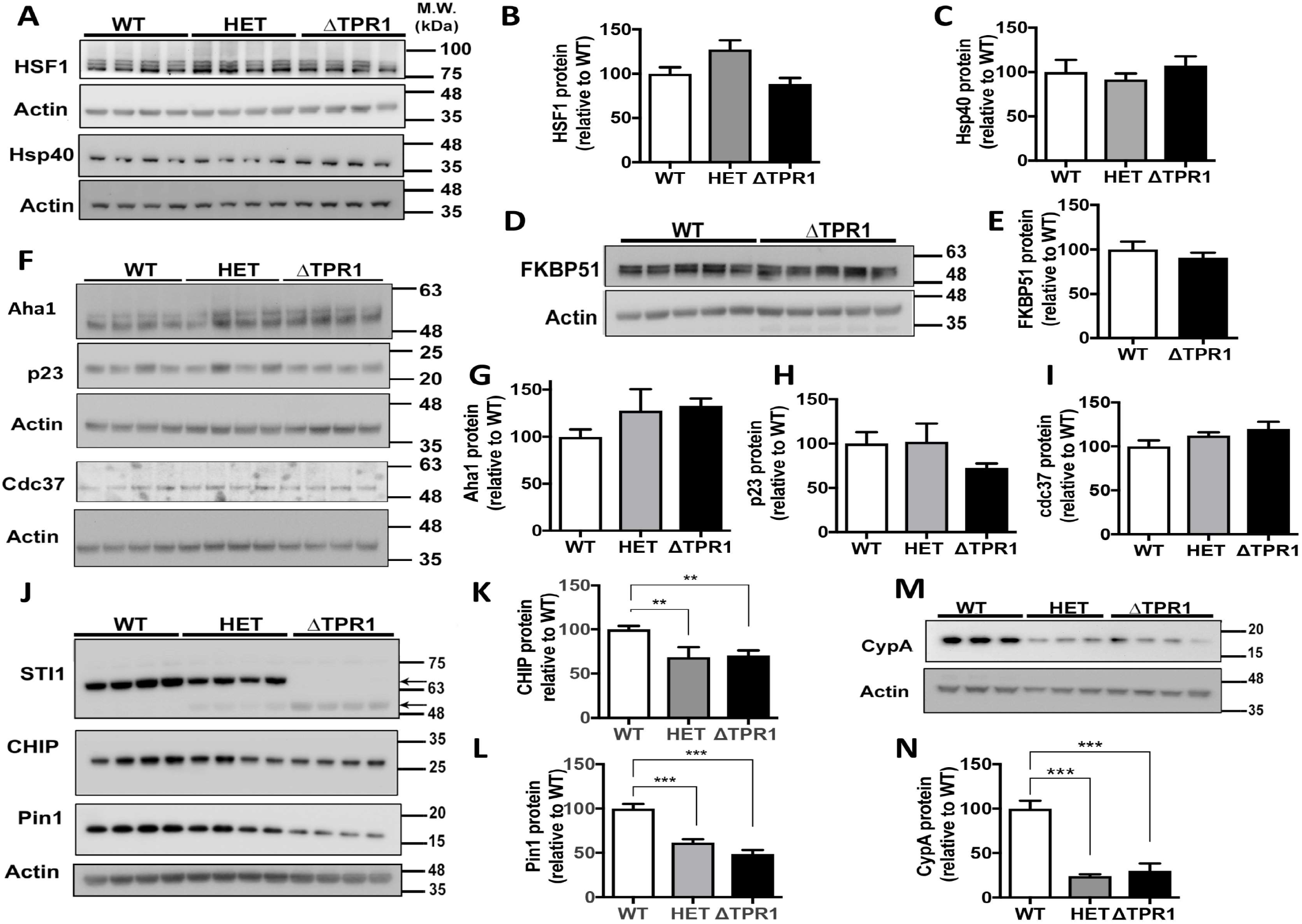
HSF1 and Hsp90 co-chaperones in aged ΔTPR1 cortices. **A.** Representative immunoblots for Heat Shock Factor 1 (HSF1) and Hsp40 in cortical extracts. **B**-**C.** Densitometric quantification of HSF1 and Hsp40 protein levels respectively (N = 4/genotype). **D.** Representative immunoblot for FKBP51 in WT and ΔTPR1 male adult cortical extracts. **E.** Quantification of FKBP 51 protein levels (N = 5/genotype). **F.** Representative immunoblots of Hsp90 co-chaperones Aha1, p23, and Cdc37. **G-I** Densitometric quantification of Aha1, p23 and Cdc37 protein levels respectively (N = 4/genotype). WT (WT/WT), HET (WT/ΔTPR1), ΔTPR1 (ΔTPR1/ΔTPR1). **J.** Immunoblot of STI1, CHIP and Pin1 in cortical lysates. **K. & L.** Densitometric quantification of CHIP and Pin1. **M. & N.** Cyclophilin A (CypA) protein expression in cortical lysates and quantification. (N = 4 for WT and ΔTPR1 and N = 4 for HET for CHIP and N = 8 for all genotypes for Pin1). All data are Mean ± S.E.M. *p < 0.05, **p < 0.01, *** p < 0.0001.

We extended our investigation to a number of proteins involved in Hsp70-Hsp90 network function or protein folding. The co-chaperone C-terminal Hsp70 Binding protein (CHIP), a ubiquitin E3 ligase that targets clients for degradation, showed 40% reduction in heterozygous and ΔTPR1 mouse brain compared to littermate controls (Fig. 4J and K, p = 0.0014). We also investigated Pin1, a peptidyl-prolyl cis/trans isomerase (PPIase) that works with the Hsp90 complex and is important for regulating tau phosphorylation (Dickey et al., 2007). In ΔTPR1 mice, Pin1 showed 50% reduction in both heterozygous and ΔTPR1 brains compared to littermate controls (Fig. 4J, L, p < 0.0001). Cyclophilin A (CypA), another PPIase, presented 75% reduction in protein levels in heterozygous and ΔTPR1 brains (Fig. 4M, N, p = 0.0004). We found no significant changes in mRNA expression for any of these genes in the brain of ΔTPR1 mice (Mean ± SEM for Fkbp5: WT 1.1 ± 0.24 and ΔTPR1 0.71 ± 0.12, p = 0.26; CHIP: WT 1.0 ± 0.10 and ΔTPR1 0.77 ± 0.07, p = 0.08; Pin1: WT 1.0 ± 0.16 and ΔTPR1 0.77 ± 0.06, p = 0.19; CypA: WT 1.0 ± 0.08 and ΔTPR1 0.78 ± 0.08, p = 0.11), suggesting the possibility of disturbed proteostasis. Unexpectedly, these results indicate that STI1 is an important regulator of the stability of a group of Hsp90 regulators and co-chaperones involved in the Hsp70-Hsp90 chaperone network. Strikingly, the effect observed was as pronounced in heterozygous mice, that express approximately 50% of WT-STI1 plus 10% ΔTPR1-STI1 (compared to controls), as it was in ΔTPR1 mice that express only 20% of ΔTPR1-STI1.

As abnormal STI1 activity could alter gene expression (Gangaraju et al., 2011; Karam et al., 2017; Sawarkar, Sievers, & Paro, 2012), we performed unbiased RNA-sequencing analysis, to test whether general transcriptome changes in mutant mice could contribute to brain phenotypes. Long RNA-sequencing was performed on 5 cortical samples from STI1 homozygous ΔTPR1 mice and 5 wild-type littermate controls. RIN values for these samples ranged between 8.2 and 8.7. Illumina sequencing yielded an average of 797,963 reads per sample, with 93.69% reads mapping rate to the mouse genome. We confirmed the complete absence of STI1 exons 2 and 3 on ΔTPR1 samples compared to wild-type control tissues (data not shown). Principal component analysis and sample distance matrix analyses did not segregate between the two groups, indicating only minimal differences between the two genotypes at the transcriptome level (Supplementary Fig. 1A, B). Furthermore, RNA sequencing analysis did not reveal any significant changes in transcripts passing FDR correction (p < 0.05). Hence, it is unlikely that large general transcriptome changes contributed to phenotypes in ΔTPR1 mice.

### ΔTPR1-STI1 affects the levels of Hsp70/Hsp90 client proteins

To test whether the function of the Hsp70/Hsp90 chaperone network was intact in ΔTPR1 mice we investigated the levels of Hsp90 clients in 15-18-month-old brain tissue from ΔTPR1 mice. We tested for glucocorticoid receptor (GR), Tau protein and G protein-coupled receptor kinase 2 (GRK2), all of which are classical Hsp90 client proteins. Although qPCR analysis showed that mRNA levels for these three classical Hsp90 clients were not altered in the cortex of ΔTPR1 mice (unpaired t-tests, Mean ± SEM for GR: WT 1.0 ± 0.09 and ΔTPR1 0.91 ± 0.08, p = 0.41; for Tau: WT 1.0 ± 0.19 and ΔTPR1 0.86 ± 0.06, p = 0.55, for GRK2: WT 1.0 ± 0.16 and ΔTPR1 0.83 ± 0.05, p = 0.28), immunoblot analyses showed that protein levels of all of these Hsp90 client proteins were very sensitive to reduced STI1 activity. GR was greatly decreased even in heterozygous ΔTPR1 tissue (Fig. 5A and B, p < 0.0001). Immunofluorescence experiments further confirmed that GR levels were decreased in neurons, without changes in GR localization (Fig. 5C). Total Tau was also reduced by immunoblotting analysis (Fig. 5D, E, p < 0.005). Likewise, GRK2 was reduced in heterozygous and homozygous ΔTPR1 mice (Fig. 5D, F, p < 0.01). As observed for the Hsp70-Hsp90 network modulators, the deficit on the stability of the different client proteins was as pronounced in heterozygous mice as it was in ΔTPR1 mice.

**Figure 5.**
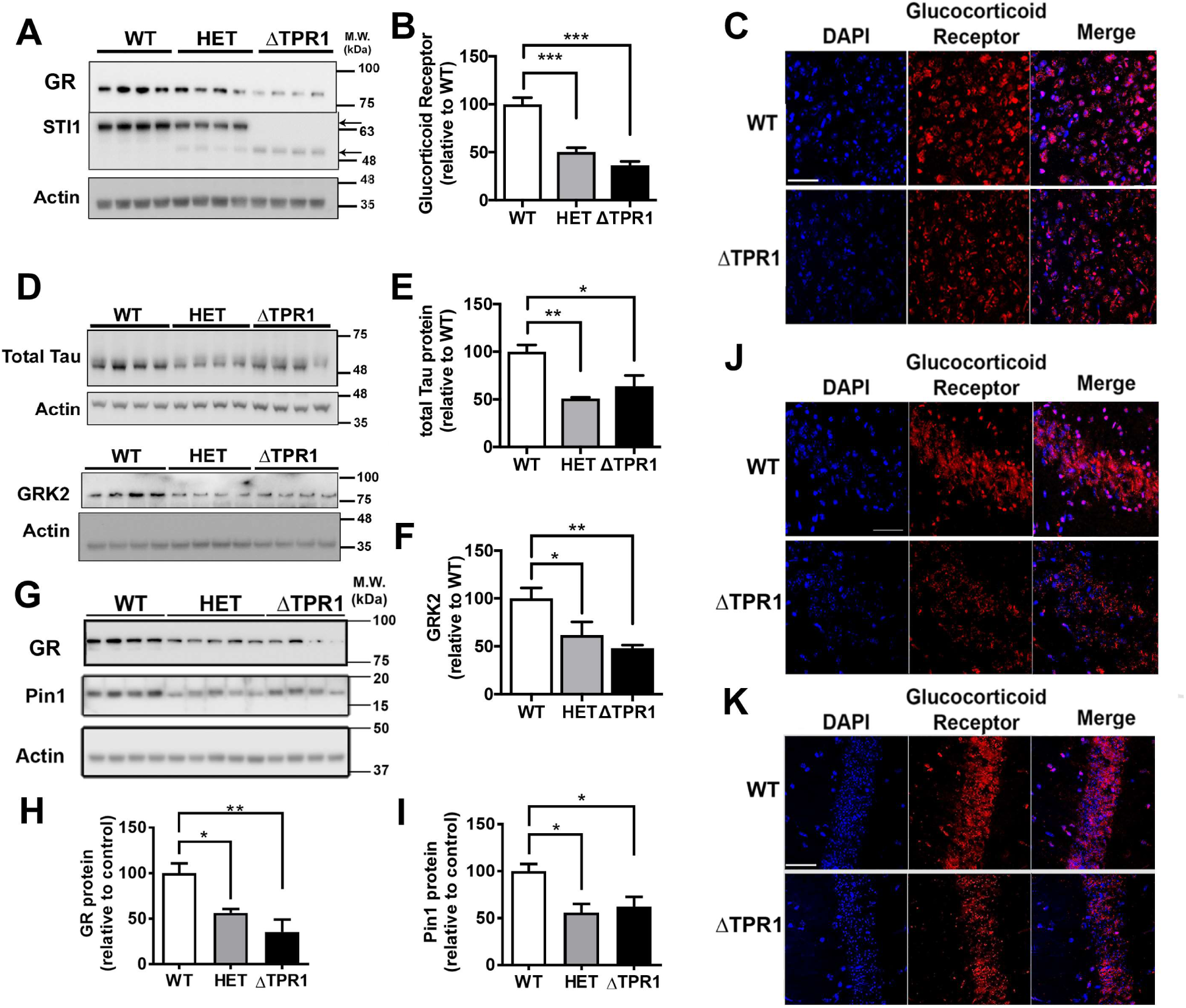
Disturbed client protein levels in aged ΔTPR1 mouse brain. **A.** Representative immunoblots for glucocorticoid receptor (GR) and STI1 in aged cortical lysates and actin loading control in male cortical lysates. **B.** Quantification of glucocorticoid receptor levels (N = 8/genotype). **C.** Confocal image of Glucocorticoid receptor expression in cortex of 15-18-month-old mice (63x). Zoom inset of 1.5. Scale bar 10 µm. **D-F.** Tau and GRK2 protein expression in cortical lysates. **F.** Quantification of total tau levels (N = 4/genotype). **F.** Quantification of GRK2 levels in cortical brain homogenates (N = 7/genotype). Data are Mean ± S.E.M. *p < 0.05, **p < 0.01, *** p < 0.0001. **G.** Representative immunoblots probing for the levels of GR and Pin1 hippocampal lysates from 15-18-month-old mice. **H. & I.** Quantification of GR and Pin1 levels (N = 4 for all blots). **J. & K.** Representative images of GR staining in CA3 and CA1 of 18-month-old WT and ΔTPR1 male mice. WT (WT/WT), HET (WT/ΔTPR1), ΔTPR1 (ΔTPR1/ΔTPR1). All data are Mean ± S.E.M. *p < 0.05, **p < 0.01, *** p < 0.0001.

We also tested whether changes in the Hsp70/Hsp90 chaperone network observed in the cortex were observed in other brain tissues by examining hippocampal samples for the level of GR (a client representative) and Pin1 (a co-chaperone representative). As observed in the cortex, immunoblot analysis of the hippocampus showed reduction in GR levels in both heterozygous and homozygous ΔTPR1 mutants (Fig. 5 G-H, p < 0.005; WT vs HET adjp = 0.016, and adjp = 0.002 for WT vs ΔTPR1 comparison). Likewise, Pin1 was also significantly reduced (Fig. 5 G and K, WT vs HET adjp = 0.013, and WT vs ΔTPR1 adjp = 0.036). Immunofluorescence analysis also showed reduction of GR in CA3 (Fig. 5J) and CA1 hippocampal neurons (Fig. 5 K) and confirmed that GR localization was not affected. These results suggest that the role of STI1 on the modulation of the Hsp70/Hsp90 chaperone network is likely identical in different neurons and brain regions.

### Hsp70-Hsp90 chaperone network dysfunction observed in ΔTPR1 mice mimics loss of STI1 function

To test whether the changes we observed in ΔTPR1 mice are reminiscent of loss of STI1 function, we generated a neuronal cell line lacking STI1, as STI1-KO embryos and STI1-KO MEFs are not viable (Beraldo et al., 2013). We used CRISPR-Cas9 technology to generate SN56-STI1-KO cells (Fig. 6) and tested them for the levels of different members of the Hsp70-Hsp90 chaperone network. Similar to what we observed for the ΔTPR1 mice, levels of Hsp90 and Hsp70 did not differ between SN56-STI1-KO and control cells (Fig. 6 A–D, panHsp90, p = 0.70; for Hsp70, p = 0.70). Likewise, levels of the client proteins GR and GRK2 were significantly decreased in SN56-STI1-KO cells when compared to control cells (Fig. 6E, F, I, GR, p = 0.0023; for GRK2, p = 0.0006). The co-chaperones CHIP (Fig. 6E, G, p = 0.0034), and CypA (Fig. 6H, K, p = 0.0017) were also significantly decreased in SN56-STI1-KO cells. Furthermore, levels of the co-chaperone FKBP51 were not altered (Fig. 6H, J, p = 0.29). Noteworthy, transfection of SN56-STI1-KO cells with STI1-HArescued the levels of GR (Fig. 6L-M, one way ANOVA, KO-HA vs KO-STI1-HA adjp = 0.0006), Pin1 (Fig. 6L and 6N, KO-HA vs KO-STI1-HA adjp = 0.0042) and CypA (Fig. 6L and 6P KO-HA vs KO-STI1-HA adjp = 0.018), further supporting the notion that the Hsp70/Hsp90 chaperone network dysfunction observed in these STI1-KO cells is dependent on STI. Interestingly, levels of CHIP were not altered by STI1-HA transfection (Fig. 6L and 6O, KO-HA vs KO-STI1-HA adjp = 0.72). These results indicate that the Hsp70-Hsp90 chaperone network dysfunction we observed in ΔTPR1 mice strongly mimics the loss of STI1, suggesting that Hsp90/Hsp70 chaperone network is highly dependent on STI1 functional levels.

**Figure 6.**
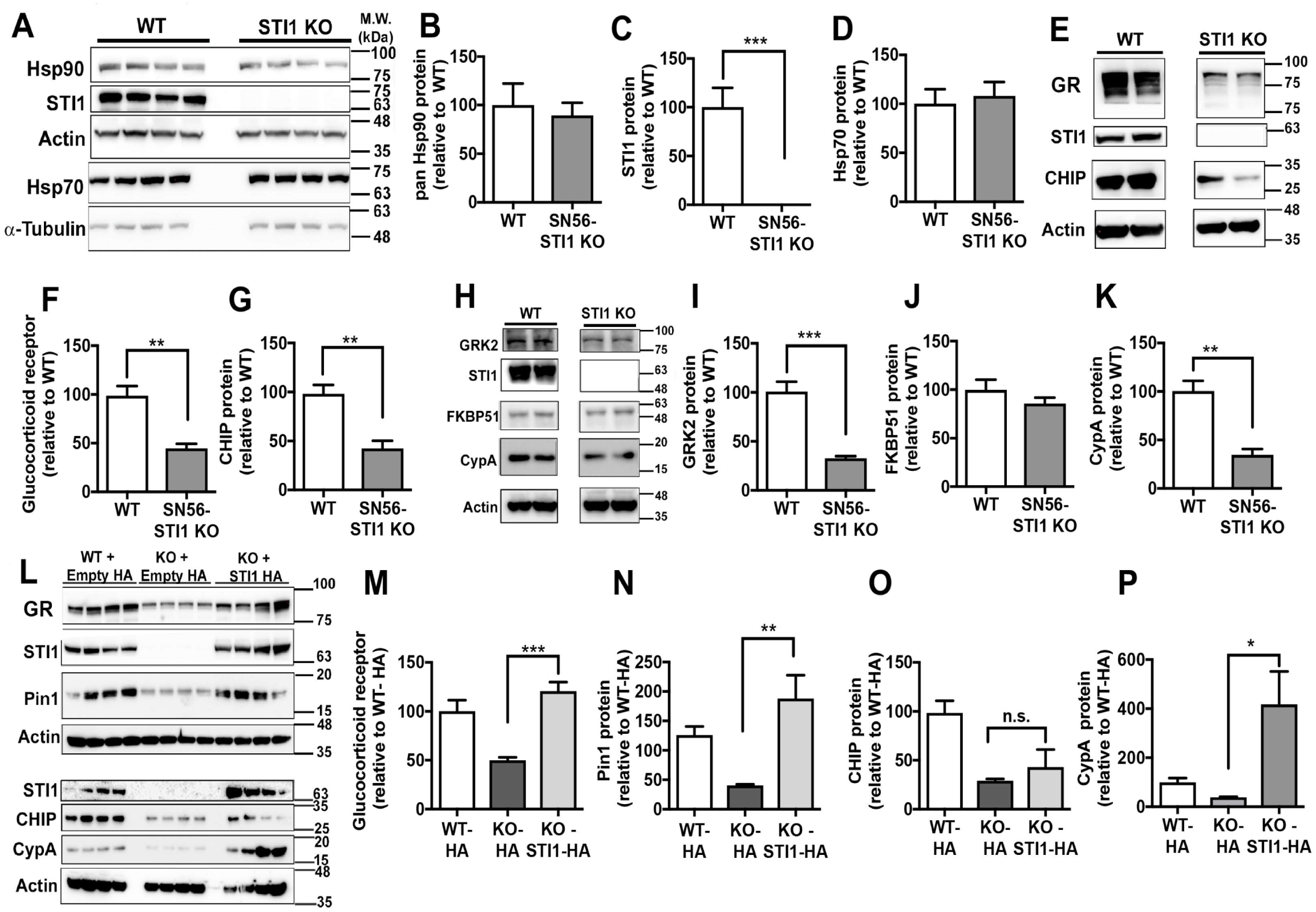
CRISPR/Cas9 SN56-STI1 KO cells have similar disruption of client and co-chaperone expression as ΔTPR1 mice. **A.** Representative immunoblots for panHsp90, STI1 and Hsp70 expression in SN56 cells. **B-D.** Densitometric quantification of Hsp90, STI1 and Hsp70 protein expression relative to WT cells. **E.** Representative **i**mmunoblots for glucocorticoid receptor, STI1 and CHIP in SN56-STI1 KO cells (images from the same blot, just cropped due to spaces left between samples) **F.** Quantification of GR levels. **G.** Quantification of CHIP levels. **H.** Representative immunoblots for GRK2, FKBP51 and CypA protein levels. **I-K.** Densitometric quantification of GRK2, FKBP51 and CypA protein levels respectively (images from the same blot, just cropped due to spaces left between samples). **L.** Rescue experiments with HA-tagged STI1. **M-P**. Quantification by densitometry for GR, Pin1, CHIP and CypA respectively. Data analyzed using Student’s t - test (N = 4 dishes)- for N-P, One-way ANOVA. STI1-KO in immunoblots represents the SN56-STI1 KO cells. All data are Mean ± S.E.M. *p < 0.05, **p < 0.01, *** p < 0.0001.

### STIP1 and co-chaperone loss of function in humans

Our experiments demonstrate that perturbation of STI1 function in mice decreases client protein levels and also impacts a number of Hsp90 regulatory proteins, some of which have not been directly linked to STI1 modulation. To determine whether similar constraint is found in humans for *STIP1* and members of the Hsp90 machinery, we determined the frequency of variation in *STIP1* in healthy individuals using public databases, combining genetic information from thousands of exomes and genomes such as ExAC (60,706 individuals) and gnomAD (138,632 individuals) (Lek et al., 2016). We observed 1 and 4 heterozygous *STIP1* protein truncating variants (PTVs) carriers in ExAC and gnomAD, respectively, at a frequency of <<0.001% (Supplementary Table 1). In comparison, *HSP90AA1* presented 8 and 22 PTVs in ExAC and gnomAD, respectively, i.e. at a 10 - fold higher frequency of <0.01% than *STIP1* (Supplementary Table 1). The *STIP1* pLI score, which reflects the probability that a given gene does not tolerate loss-of-function variation was 1, suggesting that *STIP1* loss-of-function is most likely not tolerated in humans or may result in a disease phenotype (Lek et al., 2016). In comparison, the pLI score of *HSP90AA1*, the stress inducible Hsp90 allele (α isoform), was 0.68, suggesting that PTVs may be tolerated in *HSP90AA1*. This is likely due to compensation by the highly redundant *HSP90AB1*, the constitutive Hsp90 (β) isoform. Interestingly, the *HSP90AB1* pLI score is 1. These analyses mirrored the survival of STI1, Hsp90α and Hsp90β knockout mice (Beraldo et al., 2013; Grad et al., 2010; Voss, Thomas, & Gruss, 2000). Whereas Hsp90α knockout mice survive to adulthood (Grad et al., 2010), both STI1 and Hsp90β gene ablation causes embryonic lethality (Beraldo et al., 2013; Voss et al., 2000).

We extended our analysis to other Hsp90 co-chaperones that are affected by changes in STI1 levels to determine whether they may be redundant in mammals. We did so by comparing human genetic data with viability of published knockout mice. This analysis is summarized in Table 2. Supplementary Table 1 tabulates the observed PTVs, SNVs, indels, CNVs, and associations, providing a comprehensive summary of the human genes investigated using publicly available datasets of healthy controls and disease-ascertained individuals. Taken together, our analysis suggests that constitutive Hsp90, STI1, CDC37, Aha1 and p23 are essential in mammals, indicating that some co-chaperones (such as STI1, Aha1 and p23), which are otherwise not essential in yeast (Sahasrabudhe, Rohrberg, Biebl, Rutz, & Buchner, 2017), may provide more sophisticated regulation in mammals.

**Table 2.**
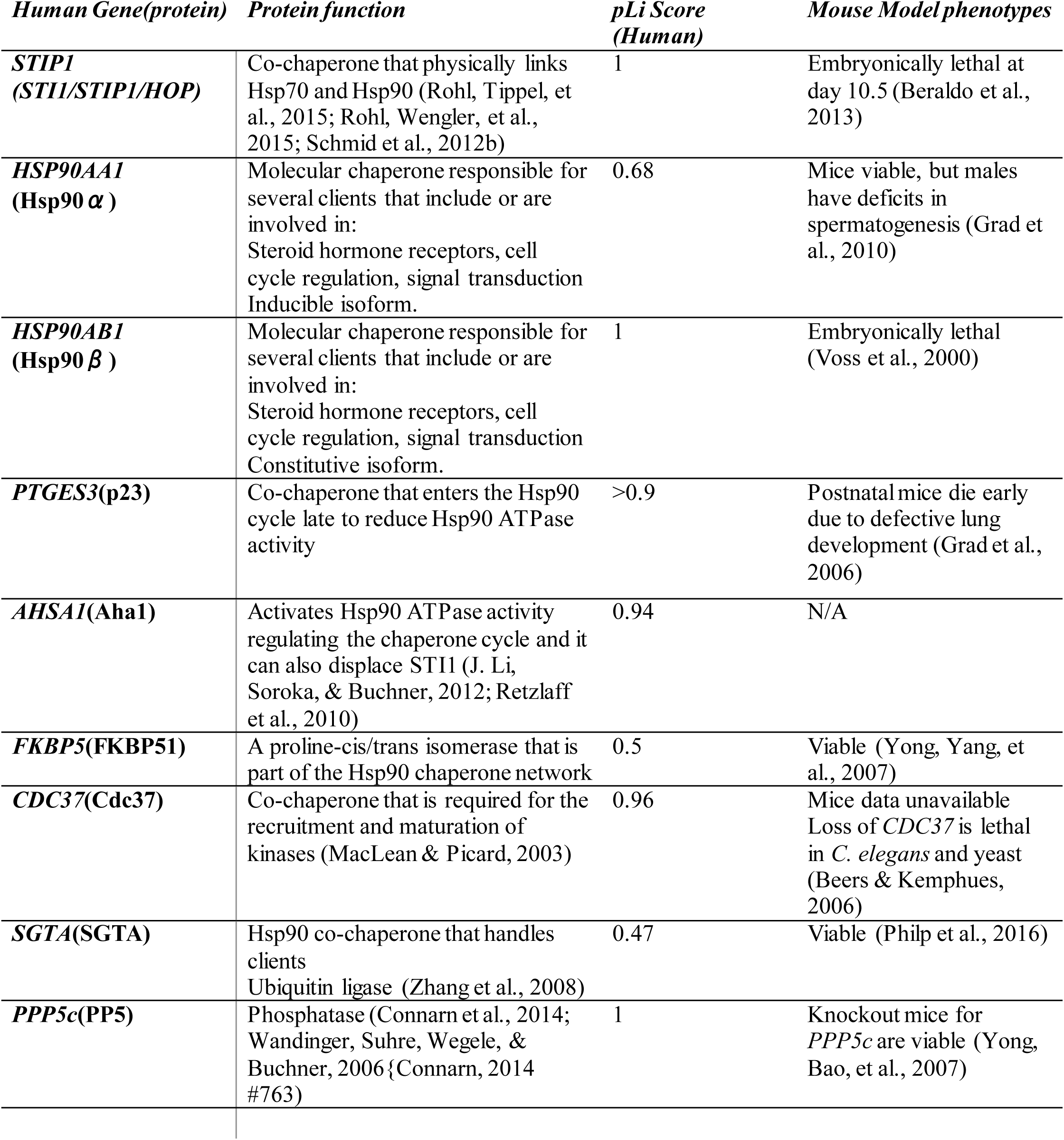
pLi score generated from human genetic data of Hsp90 co-chaperones, known function and comparison to published knock-out mouse models.

### Decreased STI1 activity compromises viability of cultured cells

Given that disturbed STI1 activity in mammalian cells interferes with different aspects of the Hsp70/Hsp90 chaperone machinery, we tested whether STI1 protects neuronal cells from environmental stress. In normal conditions, SN56-STI1 KO cells examined using the Live/Dead staining assay showed decreased survival when compared to WT controls (Fig. 7A, p < 0.0001). Moreover, SN56-STI1 KO cells presented increased sensitivity to thapsigargin, which induces ER stress (Fig. 7B, two-way ANOVA significant effect of genotype, p < .0001; significant effect of treatment, p < .0001). Similarly, ΔTPR1 MEF cultures showed decreased survival when compared to WT MEFs (Fig.7 C&D, p = 0.018). In addition, ΔTPR1 MEFs presented decreased cell proliferation when compared to WT MEFs (Fig. 7 E&F, p < 0.0005).

**Figure 7.**
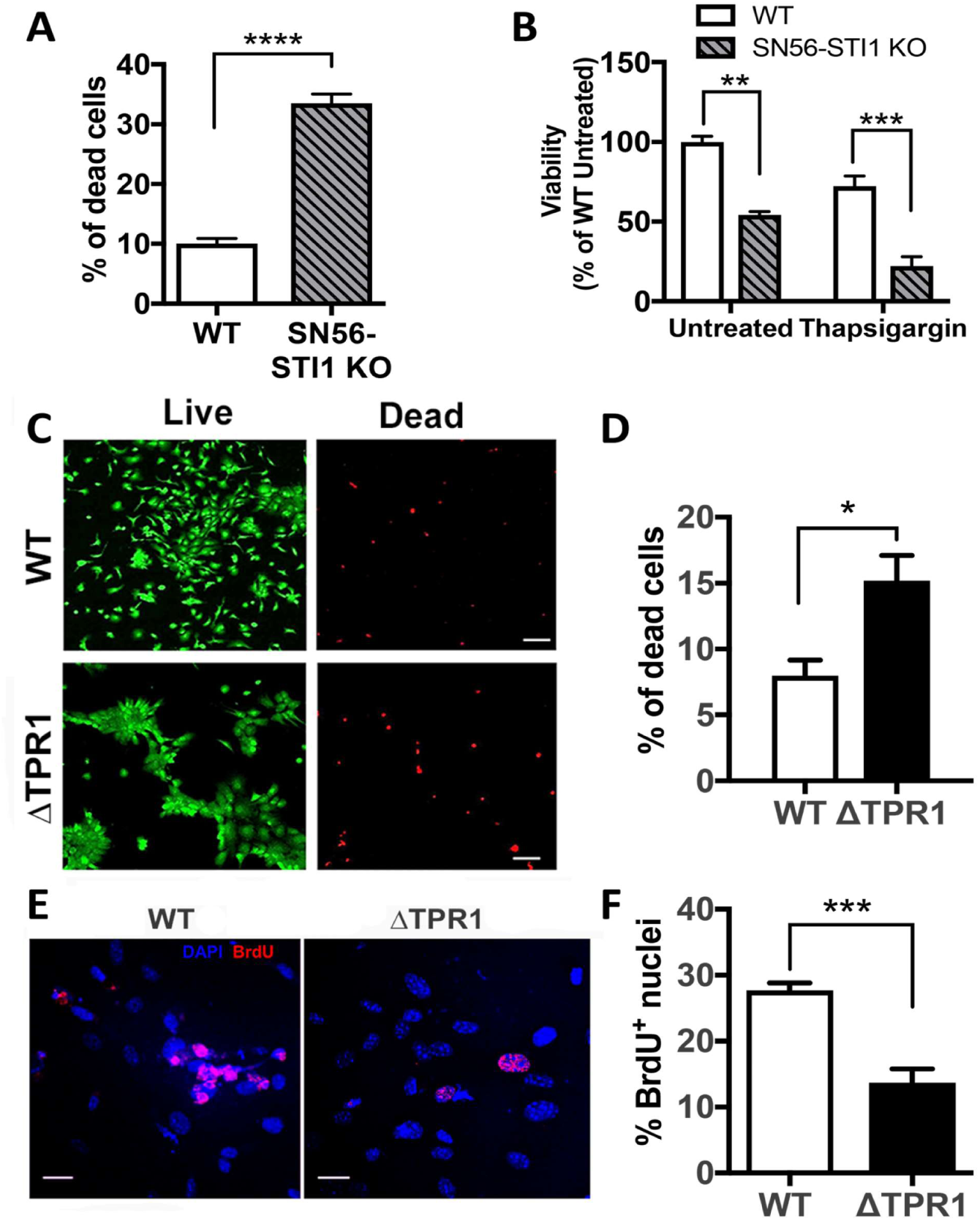
Reduced viability of cells with dysfunctional STI1. **A.** Live-Dead assay in WT and SN56-STI1 KO cells and data calculated as the percentage of dead cells (# of dead cells/#dead+#live cells). **B.** CellTiter-Glo Luminescent Cell Viability Assay in SN56-STI1 KO cells treated with 10 µM Thapsigargin in complete media for 24 h. **C.** Live-Dead Assay in P4 MEFs from ΔTPR1 mice. Scale bars 100 µm. **D.** Percentage of cell death. **E.** MEFs from ΔTPR1 mice were treated with 30 µM BrdU for 1.5 h, and then fixed and stained for BrdU. **F.** Quantification of percentage of BrdU positive nuclei/total nuclei analysis to assess MEF proliferation (Scale bars 50 µm) (N = 4-5 independent MEF cultures/genotype, Data analyzed with Student’s t-test). WT (WT/WT), ΔTPR1 (ΔTPR1/ΔTPR1). All data are Mean ± S.E.M. *p < 0.05, **p < 0.01, *** p < 0.0001.

### Impaired STI1 activity leads to age dependent decrease in the number of hippocampal neurons

Because our results showed that decreased STI1 activity compromises cellular resilience and proliferation in cultured cells, we tested whether STI1 activity is required to maintain healthy hippocampal neurons *in vivo*. We choose to study hippocampal neurons because they show increased vulnerability to a number of protein misfolding diseases (Adamowicz et al., 2017; Beyer et al., 2013; Kalaitzakis et al., 2009) and aging (Gemmell et al., 2012; Kuhn, Dickinson-Anson, & Gage, 1996; J. S. Li & Chao, 2008; Padurariu, Ciobica, Mavroudis, Fotiou, & Baloyannis, 2012). To evaluate neuronal resilience in old ΔTPR1 mice (15-18-month-old) we initially used silver staining. Silver is increasingly taken up by degenerating neurons, axons or terminals (Chen et al., 2008; Kolisnyk et al., 2016; Zhou et al., 2010). There was no difference between genotypes in silver staining in the dentate gyrus of 15-18-month-old mice (Fig. 8A and B, p = 0.12). However, we detected a significant increase in silver staining in the CA1 region of ΔTPR1 mice (Fig. 8C and D, p = 0.01). In the CA3 region of ΔTPR1 mice, silver staining showed a tendency to be increased but changes did not reach significance (Fig. 8E-H, CA3, p = 0.07). Nonetheless, CA3 sub-regions images suggested that neuronal layers (arrows) were altered in older ΔTPR1 mice, suggesting the possibility that at this age the CA3 region was already severely affected by neuronal loss.

**Figure 8.**
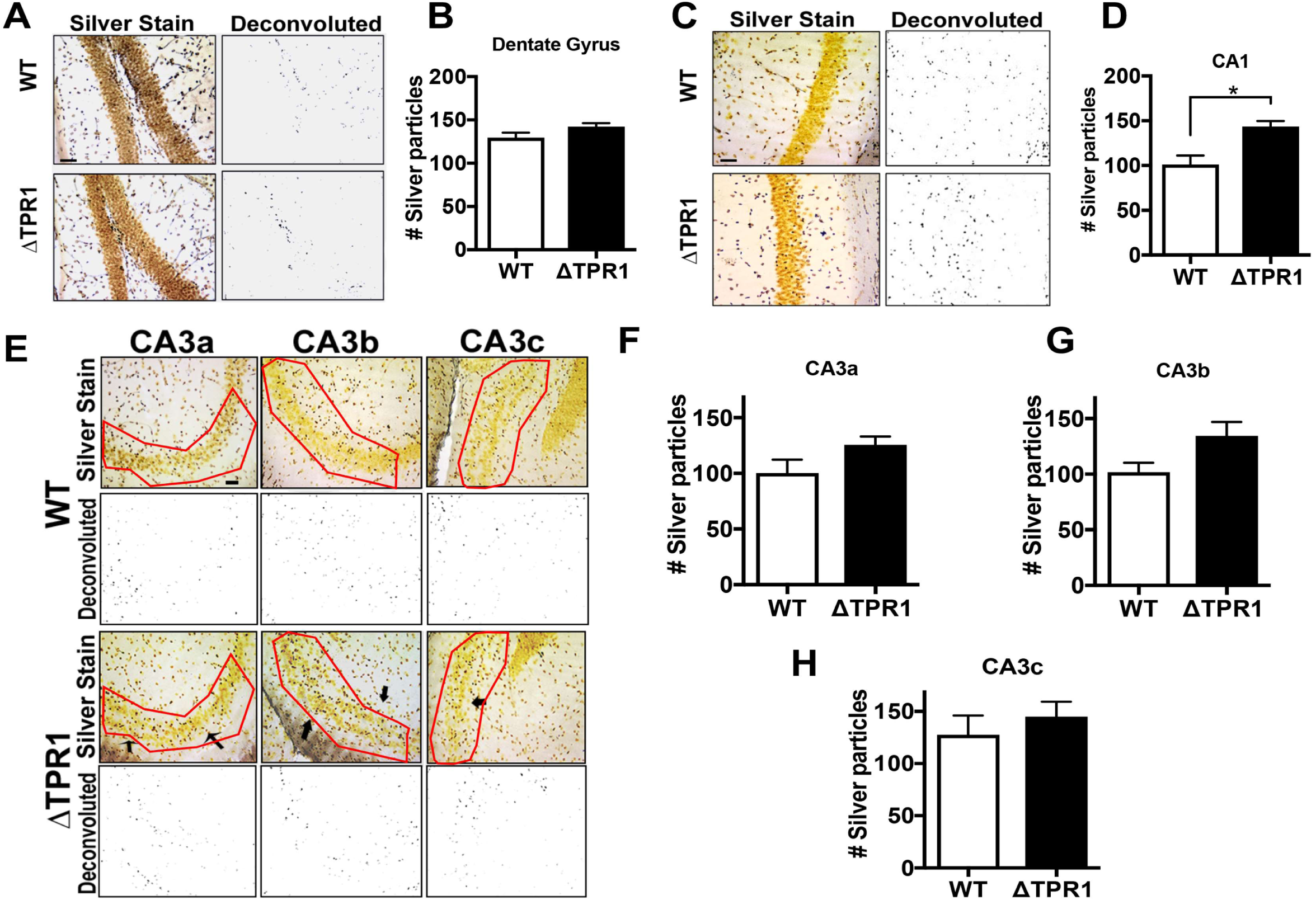
Neurodegeneration in aged ΔTPR1 mice. **A, C, E.** Silver staining in 15-18-month-old male mice. **A.** Representative images of silver staining in the dentate gyrus (at 20X magnification). Raw image and deconvoluted/thresholded image are shown to visualize silver particles. **B.** Quantitative analysis of silver particles in dentate gyrus (N = 4/genotype). **C.** Silver staining in the CA1 region. **D.** Quantification of silver staining in the CA1 region. **E.** Silver staining in CA3 subfields, CA3a, CA3b and CA3c. Arrows indicate noticeable thinning of CA3 neuronal layer in ΔTPR1 mice compared to littermate controls. **F-I.** Quantitative analysis of silver particles in hippocampal CA3 region. WT (WT/WT), ΔTPR1 (ΔTPR1/ΔTPR1). All data are Mean ± S.E.M. *p < 0.05 Student’s t-test. Scale bars 50 µm.

To further investigate neuronal survival/viability in the CA3 region of ΔTPR1 mice, we stained the hippocampus of control and ΔTPR1 mice with NeuN, a marker of mature neurons, and counted neurons in different sub-regions. Young (3-5 month) ΔTPR1 mice showed no difference in the number of CA3 neurons across all subfields when compared to WT controls (Fig. 9A-D, CA3a p = 0.22; for CA3b p = 0.22, CA3c p = 0.15). Likewise, no differences were found in CA1 region at young age (data not shown). In contrast, at 15-18 months of age, thinning of the CA3 region was obvious. Across the whole CA3, there were significantly less neurons in ΔTPR1 mutants compared to controls (Fig. 9E-H, CA3a p = 0.01; for CA3b p = 0.005; for p = 0.002). Interestingly, we compared STI1 levels in the hippocampus of 4 and 15-month-old control WT mice and observed a significant decrease with age (Fig. 9 I-J, p = 0.025). Likewise, the CA1 region had significant reduction in neuron count (data not shown, p = 0.002). As levels of the hypomorphic ΔTPR1-STI1 protein are already low in ΔTPR1 mice at young age, a further decrease may augment the stress to the chaperone system in hippocampal neurons, promoting degeneration.

**Figure 9.**
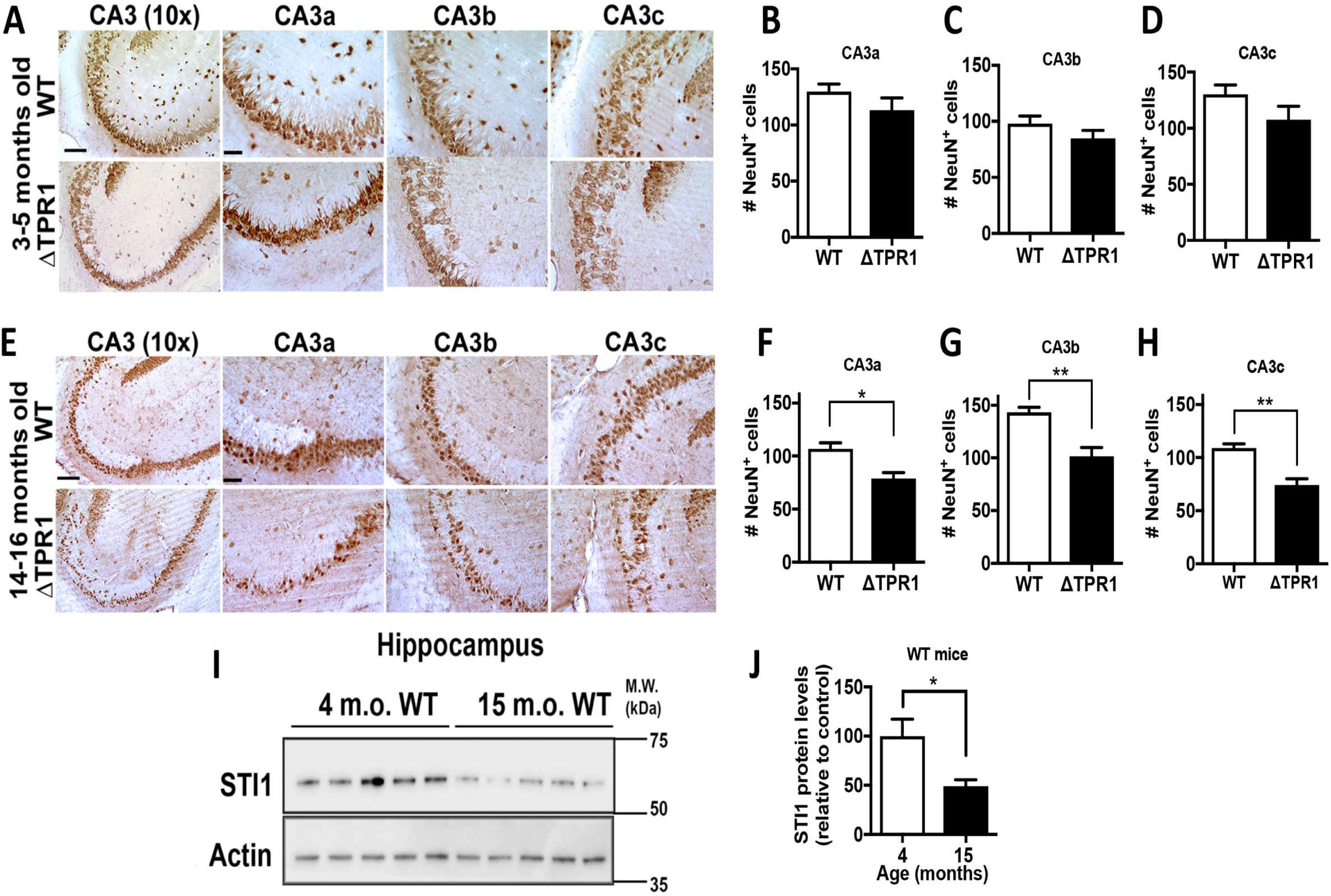
Age-dependent neuronal loss in CA3 region of the hippocampus. **A.** Representative micrographs of NeuN staining in CA3 region of 3-5-month-old male mice (at 10X magnification). **B-D.** Average NeuN positive neurons in all CA3 subfields per section. (N = 4-5 animals/genotype). **E.** NeuN staining in the CA3 region of 15-18-month-old mice (at 10X magnification). **F-H.** Average NeuN positive neurons per section relative to WT. Data analyzed using Student’s t-test (N = 4-5/genotype). **I.** STI1 expression in hippocampal lysates from C57BL/6 male mice at 4 months and 15 months of age. **J.** Densitometric quantification of STI1 protein levels, normalized to control and relative to actin. WT (WT/WT), ΔTPR1 (ΔTPR1/ΔTPR1). All data are Mean ± S.E.M. *p < 0.05, **p < 0.01, *** p < 0.0001. Scale bars represent 50 µ m for 20X images and 150 µm for 10X images.

### Spatial memory recall deficits in ΔTPR1 mice

Numerous studies implicate the CA3 region in spatial memory (Farovik, Dupont, & Eichenbaum, 2010; I. Lee, Jerman, & Kesner, 2005; J. S. Li & Chao, 2008; Steffenach, Sloviter, Moser, & Moser, 2002). To determine the functional consequence of the age dependent degeneration of hippocampal neurons, we measured performance of old ΔTPR1 mice in the spatial version of the Morris water maze (MWM). No difference between genotypes was observed in the learning phase of the task over the four days of training: on each day of training animals took less time to reach the platform (Fig. 10A, p < 0.001). On the other hand, animals from both genotypes took similar time to reach the target each day (p = 0.30). Also, on each day of training, mice swam a shorter distance to reach the platform (Fig. 10B, p < 0.001), and both genotypes swam a similar distance each day to reach the platform (p = 0.88). Interestingly, the speed of ΔTPR1 mice was slightly but significantly lower than that of WT controls (Fig. 10C, p = 0.02). These results indicate that both genotypes were able to learn the MWM task. However, on the probe trial day (fifth day), ΔTPR1 mice showed no preference for the target quadrant of the pool (Fig.10D), while control mice spent significantly more time in the target quadrant than in the other quadrants. These results indicate that while control mice clearly remembered where the platform should be, ΔTPR1 mice did not seem to retrieve this information. In summary, we observed significant degeneration of CA1 and profound loss of CA3 neurons in ΔTPR1 mice that was accompanied by a selective deficit in spatial memory.

**Figure 10.**
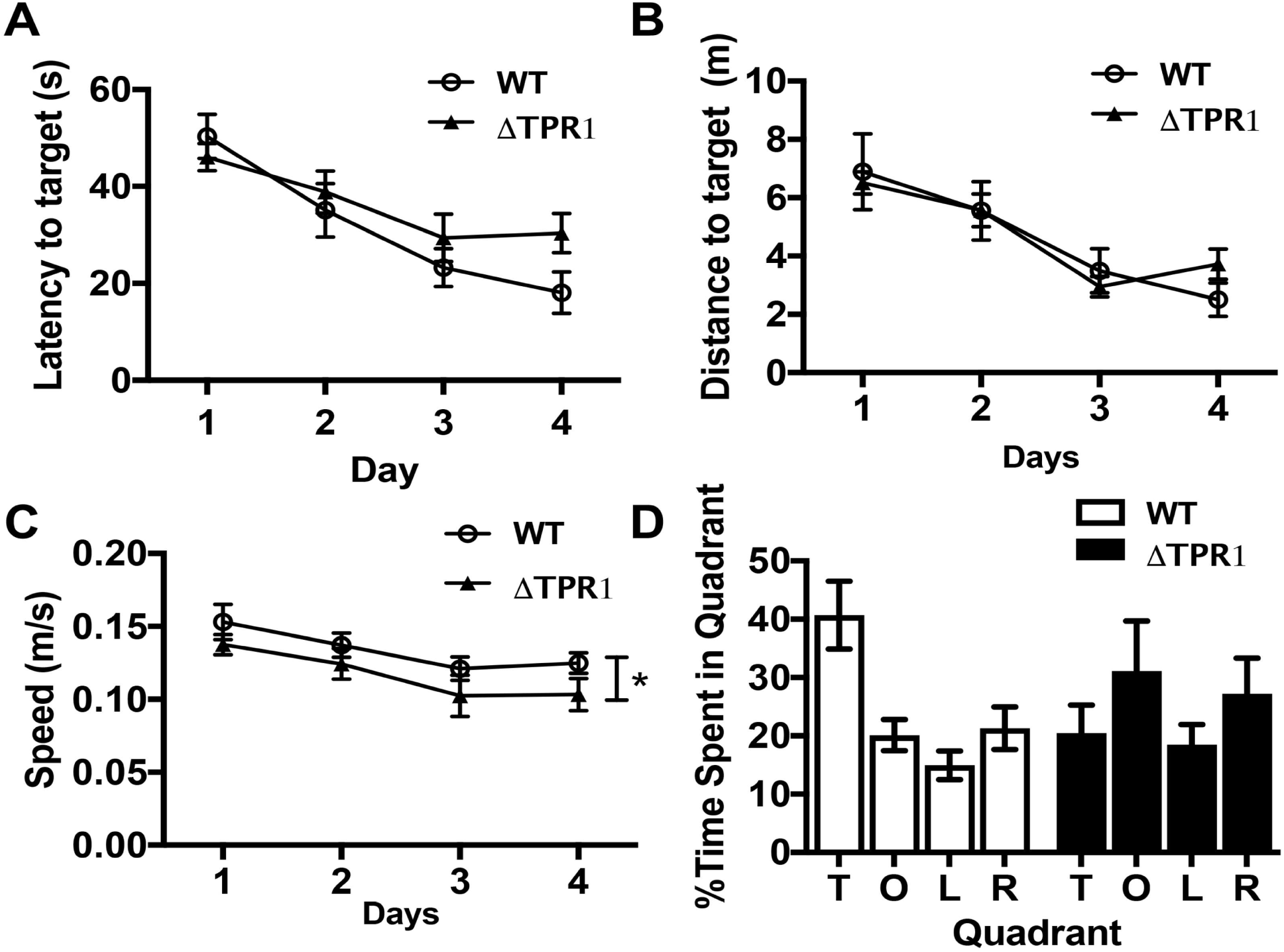
Spatial memory deficits in ΔTPR1 mice. Spatial Morris Water Maze test in 9-month-old mice. **A-C.** Measures of learning during the acquisition phase of the task. This is measured by latency to find the target platform **A.**, average distance travelled to reach platform **B.**, and mean speed (m/s) travelled before reaching platform **C.**. **D.** Represents the probe trial in which the mouse is placed in pool without platform and percentage of time spent in each quadrant is recorded. (T = target quadrant, O = opposite quadrant, L = left, R = right- with respect to target quadrant). Data analyzed using Two-Way ANOVA Repeat Measures. (n = 9-10/genotype).

## Discussion

STI1 is a highly conserved co-chaperone that plays a critical role in mediating interactions between Hsp70 and Hsp90 in the chaperone network. By taking advantage of a new hypomorphic STI1 allele in mice to understand the consequences of decreased STI1 activity/expression in the mammalian brain *in vivo*, we reveal that STI1 is a master controller of the chaperone network required to maintain stability of Hsp90 client proteins and several Hsp90 auxiliary co-chaperones in the mammalian brain. In depth RNA-Seq data and analysis of client proteins support the notion that STI1 activity regulates client levels by proteostasis, rather than by transcriptome modulation. Although the interpretation of these data needs to account for both, reduced levels of STI1 and the deletion of the TPR1 domain, the similarities between changes in clients, co-chaperones and decreased resilience between the STI1-KO SN56 cells and the ΔTPR1 mouse line, suggest that the results from this mouse line are likely due to overall decreased STI1 function.

Our analyses indicate that, in the mouse brain, proper function of the Hsp70/Hsp90 chaperone network is highly dependent on STI1 expression levels and that there is an interaction between STI1 levels and aging. The requirement for high levels of functional STI1 may be linked to the high levels of Hsp90, which accounts for 1-2% of total cellular protein in unstressed mammalian cells. Interestingly, during stress conditions, when Hsp90 levels can rise up to 4% of total cellular protein, STI1 is one of the few co-chaperones noticeably induced (Nicolet & Craig, 1989). In addition, increased levels of STI1 in a BAC transgenic mouse line is linked to augmented Hsp90 levels (Beraldo et al., 2013).

Deletion of STI1 in yeast has been shown to affect Hsp90 clients, including GR activity (but not levels) and conformation of v-Src kinase (Sahasrabudhe et al., 2017). On the other hand, other Hsp90 clients were not affected by elimination of STI1 in yeast (Sahasrabudhe et al., 2017). In contrast, in the mouse brain we found that the levels of a number of known Hsp90 client proteins, such as GR, Tau, and GRK2 were all dependent on STI1. These differences in client specificity between mammalian and yeast STI1 highlights the increased dependence of mammalian cells on regulation by co-chaperones. Our results also revealed that stability of accessory proteins with PPIase activity as well as stability of the E3 ligase CHIP are significantly reduced in ΔTPR1 mouse brain and in SN56-STI1 KO cells. Additionally, our human dataset analyses revealed that indeed co-chaperones such as p23 and STI1, which are not required for yeast survival, are essential in humans. Thus, our analysis provides, to the best of our knowledge, one of the first in depth surveys of tolerability for loss of function for different co-chaperones in mammals.

Chaperone networks have been shown to be dysregulated with age, and changes in chaperone levels in *C. elegans* can affect phenotypes due to protein misfolding (Brehme et al., 2014; H. O. Song et al., 2009; Y. Song & Masison, 2005). In *C. elegans*, KO of STI1 reduces lifespan (H. O. Song et al., 2009) and increases toxicity of Alzheimer’s related proteins (Brehme et al., 2014). Additionally, sequestration of chaperones has been observed in neurodegenerative diseases in which α-synuclein accumulates (Ebrahimi-Fakhari, Saidi, & Wahlster, 2013) and a recent transcriptome analysis revealed STI1 as one of many genes dysregulated in some rapidly progressing Lewy body dementia patients (Santpere et al., 2018). This sequestration of chaperones and co-chaperones could ultimately impair their ability to guide protein folding and maturation, thereby increasing protein aggregation, toxicity and neurodegeneration.

The hippocampus is one brain region particularly vulnerable to environmental stress, protein aggregation and neurodegeneration (Padurariu et al., 2012; Robitsek, Ratner, Stewart, Eichenbaum, & Farb, 2015; Steffenach et al., 2002). Hippocampal CA3 neurons support spatial memory (Gilbert & Brushfield, 2009) due to their excitatory and modifiable connections with the dentate gyrus and CA1 regions. Damage to the CA3 region has been shown to produce deficits in spatial memory (I. Lee et al., 2005; J. S. Li & Chao, 2008; Steffenach et al., 2002). We found that hippocampal neurons in the CA1 region show increased features related to degeneration. Most remarkably, we found a pronounced age-dependent loss of CA3 neurons. In agreement with loss of CA3 neurons we found that spatial memory recall in ΔTPR1 mice is severely compromised. The profound and widespread alteration in Hsp90 client proteins, including GR and other critical proteins involved in neuronal resilience/function, may be critical for the phenotypes observed in ΔTPR1 mice. However, whether these effects of STI1 in neurons are cell autonomous or non-cell autonomous will need to be further investigated (Lackie et al., 2017). Future experiments using conditional approaches to mutate STI1 in neurons or glia are warranted to explore these mechanisms.

In complement with our findings of decreased resilience in aging cells with compromised Hsp70/Hsp90/STI1, recent experiments have shown that partial inhibition of the Hsp90 system can increase life span in a mouse model of aging, by killing cells with a senescent phenotype that contribute to overall organism inflammation and cellular stress (Fuhrmann-Stroissnigg et al., 2017). Our experiments demonstrated that STI1 might be exploited as a key inhibitor of the Hsp90 system to influence a host of client proteins in the mammalian brain and could be used to modulate Hsp90 activity efficiently. Overall, it will be critical to find a balance between chaperone network activity that allows neurons to cope with increased stress of aging and still allow for proper disposal of damaged or senescent cells, which may be particularly reliant on Hsp90 activity to maintain proteostasis for survival (Rodina et al., 2016).

## CONCLUSIONS

Our results illuminate a requirement for optimal STI1 activity to maintain healthy hippocampal aging in mammals. Mechanistically, reduced STI1 levels can affect the efficient transfer of clients between Hsp70/Hsp90, reducing their stability, but it can also affect signalling in neurons. Our results significantly extend the knowledge about STI1 functions, centering this protein as a master regulator of chaperone activity, having an essential role for proteostasis of Hsp70/Hsp90/STI1 client proteins and survival of selective neuronal populations.

## Material and Methods

### Mouse line generation

We used Cre/loxP technology to generate mice expressing the hypomorphic *Stip1* allele lacking the TPR1 domain (ΔTPR1). Genetically-modified mice were generated by Ozgene (Perth, Australia) on a C57BL/6J ES genetic background using standard homologous recombination techniques. In short, an FRT - flanked PGK-neomycin cassette was inserted upstream of exon 2. LoxP sites were inserted upstream of the selection cassette and downstream of exon 3. The construct was electroporated into embryonic stem (ES) cells from C57BL/6J mice and targeted ES cells were injected into C57BL/6J blastocysts. Chimeric mice obtained were crossed to C57BL/6J mice to generate STI1-flox mice. To remove the selection cassette, STI1-flox mice were crossed to OzFlpE, a knock-in line that contains the FlpE variant of the *Saccharomyces cerevisiae* FLP 1 recombinase at the Rosa26 locus. Mice that had the selection cassette deleted were backcrossed to C57BL/6J to remove FlpE. The ΔTPR1 mice were obtained by crossing STI1-flox to OzCre mice (PGK-Cre at the Rosa26 locus), which allowed for germline deletion of exons 2 and 3. Backcrossing to C57BL/6J allowed the removal of the Cre transgene. Male mice were used for all experiments.

### Ethics Statement

Animals were housed and maintained at The University of Western Ontario by the Animal Care and Veterinary Services. Animals were used as outlined in our Animal Use Protocols (2016-103; 2016-104), which adhered to the Canadian Council of Animal Care (CCAC) guidelines. Animals were housed with 3-4 littermates/cage, and had ad libitum access to food (Harlan, Indianapolis, IN, USA) and water in standard plexiglass cages in a room with light/dark cycle from 7am-7pm in temperature and humidity-controlled rooms (22-25°C, with 40-60% humidity). Animals were regularly monitored by Animal Care and Veterinary Services Staff and by the researchers and technicians in the lab.

### Mouse embryonic fibroblast (MEF) culture

MEF cultures were generated as previously described (Beraldo et al., 2013; Migliorini et al., 2002). Heterozygous breeding pairs were used and E13.5 embryos were collected and isolated for culture, with 3-5 embryos/genotype being collected for each experiment. Embryos were dissected in Hanks Balanced Salt Solution on ice. Head and liver were excluded, and all other tissues were used to generate MEF cultures. Cultures were grown in 10% FBS (Gibco, Waltham, MA, USA), 1% L-Glutamine (Gibco, Waltham, MA, USA), 1% penicillin-streptomycin (10,000 U/mL, Gibco, Waltham, MA, USA) in DMEM (Wisent, St. Bruno, QC, CA). Media was changed every 3-4 days or as required. MEFs were grown for several passages and frozen at passage 2 (P2), P3, P4 and P6. Western blotting and q-PCR were performed on P4 MEFs, to guarantee that maternal STI1 was not affecting growth and patterns of protein expression (Beraldo et al., 2013). 2 × 10^6^ cells/mL were frozen in 10% DMSO, 20% FBS in DMEM for 24 hours at −80°C, then transferred to liquid nitrogen for long-term storage. Before their use for experiments, MEFs were thawed and plated in T25 flasks in media with 20% FBS, allowed to reach 70-80% confluency, then split to smaller plates in normal medium, as required.

### Generation of SN56-STI1 KO cells using CRISPR/Cas9

The guide RNAs for the mouse *Stip1* gene (STI1 Top 1: 5’CACCGGTAGTCTCCTTTCTTGGCGT 3’ and STI1 Bottom 1 5’AAACACGCCAAGAAAGGAGACTACC 3’) were designed using Optimized CRISPR Design (http://crispr.mit.edu/). They were phosphorylated, annealed and cloned at BbsI enzyme restriction site into the px330 modified vector (Addgene, Watertown, MA, USA) (Etoc et al., 2016), according to instructions from Addgene. The construct was sequenced and used to transfect SN56 cells with Lipofectamine 2000 (Invitrogen, Carlsbad, CA, USA). Clones were then isolated by serial dilution. Isolated clones were grown separately. Immunoblot analysis was used to determine clones showing complete STI1 KO. Although several clones were obtained with decreased levels of STI1, only one clone showed complete elimination of STI1 protein expression and this clone was expanded and used to further investigate Hsp90 client proteins and co-chaperones.

### Quantitative RT-PCR

RNA was isolated using the Aurum Total RNA Fatty and Fibrous Tissue Pack (Cat# 732-6870) Bio-Rad kit according to the manufacturer’s instructions. cDNA was synthesized using 2 μg RNA according to protocol (Applied Biosystems, Foster City, CA, USA). DNA was diluted and qPCR performed using SYBR Green, on a Bio-Rad CFX96 thermocycler.

Cortices from 14-16 months old perfused male mice or lysates collected from MEFs were stored in TRIzol and then frozen on dry ice before transfer to −80°C. Samples were homogenized in TRIzol and RNA was isolated using the Aurum Total RNA for fatty and fibrous tissue kit. cDNA was generated as described in Beraldo et al. (2013). β-actin was used to normalize mRNA levels and negative controls were included (four to five distinct tissue extracts per genotype on each plate). The following primers were used to assess mRNA levels. STI1-F: 5’-GCCAAGAAAGGAGACTACCAG-3’ and STI1-R: 5’-TCATAGGTTCGTTTGGCTTCC-3’for exons 2 and 3 and STI1-97F: ACCCCAGATGTGCTCAAGAA and STI1-97R: TCTCCTCCAAAGCCAAGTCA for exons 8 and 9. Hsp90α-F:CCACCCTGCTCTGTACTACT; Hsp90α-R:CCAGGGCA TCTGAAGCATTA; Hsp90β-F: CTCGGCTTTCCCGTCAAGAT, Hsp90β-R: GTCCAGGGCATCTGAAGCAT, Hsp70-R: ACCTTGACAGTAATCGGTGC, Hsp70-F: CTCCCGGTGTGGTCTAGAAA, HSF1-F: GATGACACCGAGTTCCAGCA, HSF1-R: CACTCTTCAGGGTGGACACG, CHIP-F: CTTCTACCCTCAATTCCGCCT, CHIP-R: CATTGAGAAGTGGCCTTCCGA, Pin1-177F: AAGCAGACGCTCCATACCTG, Pin1-177R: AGAGTCTGGACACGTGGGTA, Fkbp5-83F: CTGCTGTGGTGGAAGGACAT, Fkbp5-83R: TCCCAATCGGAATGTCGTGG, Nr3c1-160F: TGTGAGTTCTCCTCCGTCCA, Nr3c1-160R: GTAATTGTGCTGTCCTTCCACTG, Mapt-200F: AACCAGTATGGCTGACCCTC, Mapt-200R: TCACGTCTTCAGCAGTTGGA, Grk2-119F: CTGCCAGAGCCCAGCATC, Grk2-119R: AGGCAGAAGTCCCGGAAAAG, Actin-F: TGGAATCCTGTGGCATCCATGA, Actin-R: AATGCCTGGGTACATGGTGGTA.

### Immunofluorescence

Immunofluorescent labelling of fixed cell cultures was conducted as previously described (Beraldo et al., 2013). For MEFs, P4 cells were used. Cells were split from T25 flasks at a density of 6 × 10^4^ cells to 24-well dishes with poly-lysine coated coverslips. Once MEFs reached 80% confluence (∼2-4 days), media was removed, coverslips were washed three times with PBS and then fixed for 20 minutes with 4% cold paraformaldehyde (PFA). After three PBS 0.5% Triton X-100 washes, cells were blocked in 0.5% Triton X-100 and 5% bovine serum albumin (BSA) in PBS for one hour at room temperature (RT) before overnight incubation with primary antibodies in 0.1% Triton X-100 and 0.1% BSA in PBS at 4°C. Cells were incubated with STI1 antibody (1:200 in-house antibody generated by Bethyl Laboratories Montgomery, USA using recombinant STI1), anti-Hsp70 (1:100, Catalog# Ab2787, Mouse mAb, Abcam, Cambridge, UK), anti-Hsp90 (1:50, Catalog# 4877, Rabbit mAb, Cell Signalling, Danvers, MA, USA). Alexa Fluor-conjugated secondary antibodies (Molecular probes) were used at 1:800 in 0.5% bovine serum albumin and 0.1% Triton X-100 in PBS. Cells were counterstained with DAPI, mounted onto slides using Immu-Mount (Thermo Scientific, Waltham, MA, USA) and imaged using Leica TCS SP8 (Leica Microsystems Inc., Ontario, Canada) confocal system (63X objective, N.A. of 1.4 and 40X objective, N.A. of 1.3). Two-three coverslips per embryo were imaged and 8 random fields of view were captured for each coverslip by a researcher blind to genotypes.

### RNA-seq analysis

For RNA sequencing, 5 WT and 5 homozygous ΔTPR1 samples were used. Briefly, tissue was homogenized in TRIzol before phase separation with chloroform. After cold centrifugation, top aqueous layer containing RNA was isolated. RNA was precipitated with 100% ethanol and pellet was collected after centrifugation. RNA pellet was washed with 85% ethanol before drying and resuspending in DEPC treated water. RNA quality was determined with RNA 2100 Bioanalyzer (Agilent, Santa Clara, CA USA), and samples with RIN values ranged between 8.2 and 8.7 were used. Sequencing-compatible poly(A)-terminated single-end libraries were generated using an RNA Library prep kit (SENSE Total RNA-Seq Library Prep Kit, 009; Lexogen, Vienna, Austria) following manufacturer’s instructions. Libraries were barcoded and sequenced on a NextSeq Series Sequencing System (HUJI Center for Genomic Technologies) using Illumina flow cell (Illumina 500 NextSeq High Output v2 Kit, FC-404-2005; Illumina, San Diego, CA, USA). The sequencing data were uploaded to the usegalaxy.org (Afgan et al., 2016) web platform for further analysis. All reads were aligned to the mouse reference genome (GRCm38/mm10) with an average 93.7% mapping (TopHat2) (Kim et al., 2013). Gene expression counts were generated using HTseq-count (Anders, Pyl, & Huber, 2015) (GRCm38/mm10) and expression analysis was performed using the Bioconductor DESeq2 (Love, Huber, & Anders, 2014) software via R platform (Team, 2017). Libraries from all samples were overall similar in depth.

### Cell death and viability assay

Cell death was assessed by the Live/Dead Viability/Cytotoxicity Kit assay for mammalian cells (Cat# L3224, Thermo Fisher Scientific – Invitrogen, Waltham, CA, USA–) as previously described (Beraldo et al., 2016; Beraldo et al., 2013; Soares et al., 2013). Briefly, SN56 cells and P4 MEFs were incubated with the calcein-AM/ethidium homodimer mix according to the manufacturer’s instructions in the original medium for 45 min and then washed 3 times with Krebs-Ringer HEPES (KRH) buffer. Images were collected using the LSM 510 META ConfoCor2 equipped with a 10x/0.3 objective. 488 nm laser was used to detect for calcein (live cells) or ethidium homodimer (dead cells). Cell death levels were quantified as the percentage of dead cells relative to the total number of cells. The numbers of live and dead cells were quantified, with 4-5 embryos per genotype and each embryo in duplicate or triplicate. Eight randomized fields of view within the well were analysed.

Viability of SN56-STI1 KO cells was also assessed using CellTiter-Glo® Luminescent Cell Viability Assay (Catalog # G7570, Promega, Madison, WI, USA) which quantifies levels of ATP, which is an indirect measure of number of cells. Experiments were conducted following manufacturer’s instructions. Briefly, cells were plated in 96-well plates, serum starved then treated to lyse cells and release ATP. A recombinant luciferase is added to the cells then relative luminescence is collected using a plate reader. For the Thapsigargin (Catalog# 586005, Millipore, Burlington, MA, USA) treatment to induce ER stress, cells were treated at a concentration of 10 µM for 24 h.

### BrdU proliferation assay

BrdU proliferation assay was performed as described previously (Beraldo et al., 2013). P4 MEFs were serum starved for 24 h before 30 µM BrdU (dissolved in sterile water) was added to serum-free culture media for 1.5 h. Media was removed and cells were fixed with cold 4% PFA for 20 minutes. Cells were washed with PBS three times then treated with 2 M HCl for 30 minutes. Acid was quickly removed and 9.1 M Sodium Borate was added to cells for 12 minutes. Cells were then washed with PBS three times, blocked for 1 h in PBS + 0.3% Triton X-100 and 5% normal goat serum, and then incubated overnight at 4°C with anti-BrdU biotin conjugate (1:100, Catalog# MAB3262B, Millipore, Burlington, MA, USA), followed by incubation with Streptavidin Alexa Fluor 488 conjugate (1:800, Catalog# S32354, Invitrogen, Waltham, MA, USA). Cells were washed with PBS then treated for 20 minutes with Hoechst. The BrdU positive nuclei were quantified and compared to total nuclei, with 4-5 embryos per genotype. Experiments were replicated three times. Experimenter was blind during image capture.

### Expression of STI1 recombinant domains in bacteria

Expression of recombinant STI1 and analysis of STI1 antibody interaction with STI1 domains was performed as previously described (Maciejewski et al., 2016).

### Western Blotting

Mice were decapitated and brains were rapidly excised. Cortex and hippocampus were dissected on ice and flash frozen on dry ice before transferred to −80°C. Protein extraction and Western blot were carried out as previously described (Beraldo et al., 2013; Guzman et al., 2011). Briefly, protein was extracted from whole cell lysates or brain tissues using ice cold RIPA lysis buffer with protease and phosphatase inhibitors. Samples were sonicated using sonic dismembrator 3 × 7 s, rocked for 20 minutes then centrifuged for 20 minutes at 10,000 g at 4°C. Supernatant was collected and used for quantifying protein concentration using the Bio-Rad DC Protein assay. 5-30 µg of protein was loaded on Bolt 4-12% Bis-Tris gradient gels. The primary antibodies used in immunoblotting were: anti-STI1 (1:5000, in-house antibody generated by Bethyl Laboratories Montgomery, USA) (Beraldo et al., 2013), anti-Hsp90 (1:1000, Catalog# 4877, Cell Signalling, Danvers, MA, USA), anti-Hsp70 (1:1000, Catalog# ab2787, Abcam, Cambridge, UK), anti-Hsp40 (1:1000, Catalog# 4868, Cell Signalling, Danvers, MA, USA) anti-CHIP (1:1000, Catalog# 2080, Cell Signalling, Danvers, MA, USA), anti-Glucocorticoid Receptor (1:1000, Catalog# 3660, Cell Signalling, Danvers, MA, USA), anti-HSF1 (1:1000, Catalog# 4356, Cell Signalling, Danvers, MA, USA), anti-Hsp90β (1:1000, Catalog# 5087, Cell Signalling, Danvers, MA, USA), anti-GRK2 (1:1000, Catalog# 3982, Cell Signalling, Danvers, MA, USA) anti-FKBP51 (1:1000, ab2901, Abcam, Cambridge, UK), anti-CypA (1:2000, Catalog# ab126738, Abcam, Cambridge, UK), anti-tau H150 (1:200, Catalog# sc-5587, Santa Cruz, Dalla, TX, USA), anti-Pin1 (1:250, Catalog# sc-15340, Santa Cruz, Dalla, TX, USA), anti-Ahsa1 (1:1000, Catalog# 12841, Cell Signalling, Danvers, MA, USA), anti-Cdc37 (1:1000, Catalog# 4793, Cell Signalling, Danvers, MA, USA), anti-p23 (1:1000, Catalog#: NB300-576, Novus Biologicals; Biotechne, Littleton, CO, USA). Protein expression was quantified using the Alpha Innotech software for the FluoroChemQ chemiluminescent exposure system (Alpha Innotech; GE Healthcare, London, ON, Canada) or ImageLab for ChemiDoc system (BioRad, Hercules, CA, USA). Expression data was relative to β-actin (1:25000, Catalog # A3854, Sigma, St. Louis, MO, USA) and normalized to WT controls. At least 2 independent blots were produced with no less than four animals analyzed for each protein (generally n = 4-9).

### Silver Staining

Silver staining was performed as described previously (Kolisnyk et al., 2016). Briefly, after trans-cardial perfusion, mouse brains were fixed in 4% PFA for 48 h before long-term storage in PBS + 0.02% sodium azide. Brains were cut using Leica VT1000S Vibratome, at 30 µm thickness and sections were stored free floating in PBS + 0.02% sodium azide. Using 6-well plates and net-wells, free-floating sections were stained with the NeuroSilver^TM^ staining kit II (Catalog#: PK301, FD NeuroTechnologies, Inc., Baltimore, MD, USA) following manufacturer’s instructions. This kit labels degenerating neuronal bodies, processes and terminals. Images were taken using Zeiss Axioskop Optical Microscope at 20X magnification, with two images being taken along the dentate gyrus, from the apex to the hilus/opening of the blades, three images of the CA3, respective to CA3 subfields (CA3a, CA3b, CA3c), and one image from the CA1 region. At least 4 sections from 4 animals/genotype were stained and sections selected were at least 120 µm apart. Using ImageJ (Fiji) Software (NIH, Bethesda, MD, USA), images were converted to 8 bits and thresholded to make the silver particles black and background white (Circularity of 0-0.65). Particles were numbered and averaged for each animal. The same parameters were used for each section, animal and both genotypes.

### NeuN Staining and Hippocampal Neuron Density

NeuN staining was performed as described previously (Kolisnyk et al., 2016). Briefly, 30 µm thick sections were mounted onto charged microscope slides, boiled in 10 mM sodium citrate for 20 minutes and then cooled for 40 minutes to room temperature for antigen retrieval. Sections were then washed three times in water, then immersed in 3% hydrogen peroxide for 10 minutes, followed by two washes in 0.025% Triton-X PBS. Tissue was blocked in 10% normal goat serum and 1% BSA in PBS for 2 h, at RT. Sections were incubated with anti-NeuN (1:15000, Catalog# ab104224, Mouse mAb, Abcam, Cambridge, UK) overnight at 4°C, washed twice with 0.025% Triton-X in PBS, then incubated for 1 h in biotinylated goat anti-mouse secondary antibody (1:200, Catalog# BA9200, Vector Laboratories, Burlingame, CA, USA). Following washes, sections were incubated for 30 minutes in Vectastain Elite ABC Kit (Catalog# PK-6100, Vector Laboratories, Burlingame, CA, USA) following manufacturer’s instructions. Next, sections were treated with DAB Peroxidase Substrate Kit (Catalog# SK-4100, Vector Laboratories, Burlingame, CA, USA) following manufacturer’s instructions. Sections were washed, then dehydrated and cleared using a series of ethanol and xylene washes. Staining was performed in 3-5 months old and 14-16 months old mice. Four sections at least 60 µm apart were selected from each animal to allow for unbiased selection within similar coordinates for each animal, starting around Lateral 2.356 mm (sagittal sections). Images were taken using Zeiss Axioskop Optical Microscope at 20X magnification. For each section, one image of each subfield of the CA3 region of the hippocampus was taken (CA3a, CA3b, CA3c) and one along the CA1. The number of NeuN cells in each subfield of the CA3 was counted, in each section and averaged per animal. The number of NeuN positive cells was then averaged across four to five animals, per genotype. By adding the number of cells across each CA3 subfield, the average sum of neurons in the CA3 was also quantified. The experimenter was blind to genotypes during imaging and quantification.

### Metabolic Cages

Analysis of mouse activity, food and water intake, oxygen consumption, carbon dioxide production and sleep cycles was analyzed using The Comprehensive Lab Animal Monitoring System as previously described (Guzman et al., 2013; Janickova et al., 2017; Roy et al., 2013).

### Morris Water Maze (MWM)

The spatial version of the MWM was performed as described elsewhere (Beraldo et al., 2015; Kolisnyk et al., 2016; Kolisnyk et al., 2013). Nine-ten animals/genotype were used in MWM based on previous estimates using this protocol (Beraldo et al., 2015; Kolisnyk et al., 2016; Kolisnyk et al., 2013) and mice were tested at 9 months of age. Animals were allowed to acclimatize to the room for 30 minutes before the start of the experiment. Experiments were performed in a 22-24°C room. Mice were placed in a 1.5 m diameter pool (water temperature was 26°C) with a transparent plastic platform 1 cm below water surface. Two large lamps next to the pool were lit all throughout the experiment and spatial cues were present on the walls. Mice were counterbalanced and trained over four days to find a platform in one quadrant of the water maze. Animals had four training sessions per day lasting 90 s each (with a 15 min inter-trial interval). If a mouse failed to reach the platform it was positioned on it for 10 seconds before being removed from the pool. In analysis, a 60 s latency to reach platform value was input for animals that exceeded 60 s before reaching the platform and ending the trial. On the fifth day, mice were placed in the pool once (without the platform) for 60 s. Time spent in target quadrant was compared to other quadrants. Activity and behaviour were recorded with ANY-Maze Software. The researcher was not blind to genotypes during experiments, and animals were randomly allocated as for the order they did the task. Target quadrant and analysis was blind.

### Analysis of Hsp90 and co-chaperone variants in human datasets

We evaluated the frequency of loss-of-function variants (protein truncating variants, PTVs) in multiple large datasets aggregating human genetic information. Specifically, we sought PTVs in the following genes: STI1, HSP90AA1, HSP90AB1, PTGES3, AHSA1, FKBP5, PIN1, STUB1, CDC37, PPIA, PPP5C, and SGTA. To assess the frequency of PTVs in these genes in a large and ethnically diverse cohort of relatively healthy individuals, and to extract the probability of loss-of-function (pLI) score for each gene, we used the Exome Aggregation Consortium (ExAC, n = 60,706) and the Genome Aggregation Database (gnomAD, n = 138,632), (Consortium, 2013; Lek et al., 2016). We also assessed the frequency of PTVs, missense variants, insertions, deletions, and/or duplications in these genes of interest in a cohort of clinically-ascertained individuals. To this end, we used the GWAS Catalog, ClinVar, and DatabasE of genomiC varIation and Phenotype in Humans using Ensembl Resources (DECIPHER) (Firth et al., 2009; Landrum et al., 2016; MacArthur et al., 2017; Mick et al., 2011; Miller et al., 2010; Stenson et al., 2017). We did not use the Human Gene Mutation Database (HGMD) as this resource is no longer open-source, and currently only provides minimal information for variants identified in patients. We applied the following criteria to ensure that PTVs observed are likely to be true PTVs: 1) ensuring the PTVis present in the canonical transcript; 2) the PTV occurs prior to the last exon of the gene; and 3) the site and the surrounding region of the PTV are sufficiently covered, 4) there is adequate allele balance between the reference allele and the alternate allele for heterozygous PTVs, and 5) the variant does not display strand bias. For variation reported by ClinVar and DECIPHER, we only included information on cases with single nucleotide variants (SNVs), small insertions or deletions (indels), or copy number variation (CNV, gain and loss) of large genomic regions, that were classified as either ‘pathogenic’ or ‘likely pathogenic’ by each database’s respective criteria. Due to the difficulty in sequencing regions of HSPA1A (Hsp70), an accurate pLI score was not obtained for this gene.

### Statistical Analyses

Data were compiled and analyzed using GraphPad Prism 6.0 Software and results are represented as Mean ± SEM. Comparison of two groups was analyzed using Student’s t-test (two sided). For groups larger than two, One or Two-Way ANOVA or Repeated Measures Two-Way ANOVA was used when needed and Post-hoc tests corrected for multiple comparisons was used when required. Mouse birth frequencies were analysed by the Chi-Square test.

**This manuscript was previously submitted as a pre-print using BioRxiv. https://doi.org/10.1101/258673**

## Acknowledgements

We thank Jose Marques-Lopes for help in generating MEFs. MAMP and VFP received support from the Canadian Institutes of Health Research (MOP 126000, MOP 136930, MOP 89919), National Science and Engineering Research Council of Canada (402524-2013 RGPIN). MAMP, MLD and FB received support from the ALS Canada. Initial generation of STI1 mutant mice was supported by PrioNet-Canada and FAPESP (Brazil) to MAMP, VFP and VRM. MHL received a FAPESP Sabbatical fellowship (2016/00440-9) and a research grant (2017/20271-0). HS acknowledges support by The Israeli Ministry of Science, Technology and Space, Grant No. 53140, The Israel I-Core Center of Excellence for Mass Trauma, the Legacy Heritage Science Initiative (LHSI) of The Israel Science Foundation Grant No. 817/13, and the Edmond and Lily Safra Center for Brain Sciences (ELSC). REL received support from OGS and the Alzheimer’s Society of Canada through the Alzheimer’s Society Research Program. SMKF is supported by the ALS Canada Tim E. Noël Postdoctoral Fellowship. The funders had no role in study design, data collection and analysis, decision to publish, or preparation of the manuscript.

## Author contributions

Designed experiments, performed experiments and analysed data: REL, AR, GM, FHB, AM, RG; Performed experiments and analysed data: JF; Analysed data: SMKF, W-YC, DG; Contributed with reagents and special tools: MHL, VRM, MLD; Designed experiments: HS, VFP, MAMP; Wrote manuscript: REL, SMKF, VFP, HS, MAMP. All authors edited and approved the final version of the manuscript.

## Figure Legends

**Supplementary Table 1:**
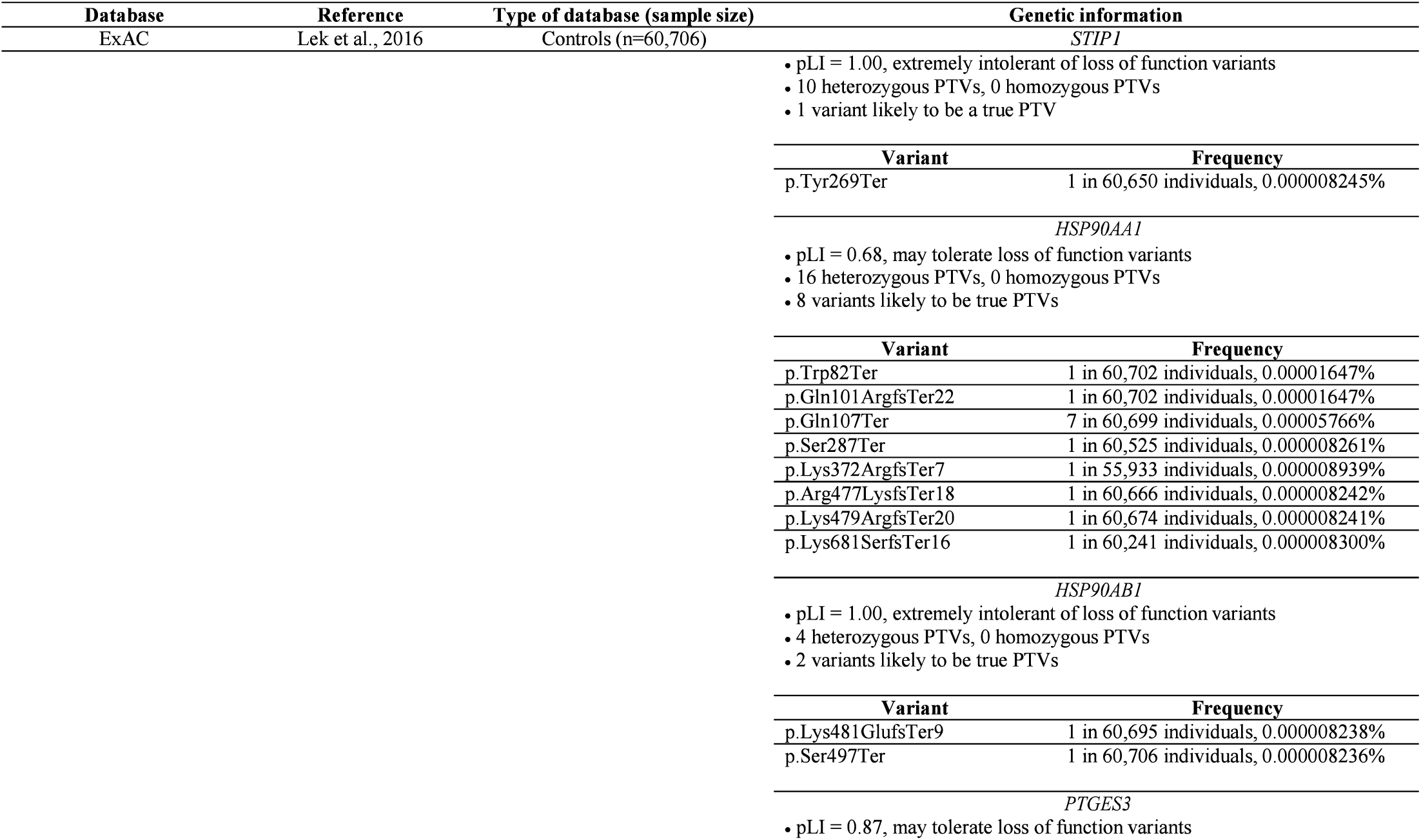
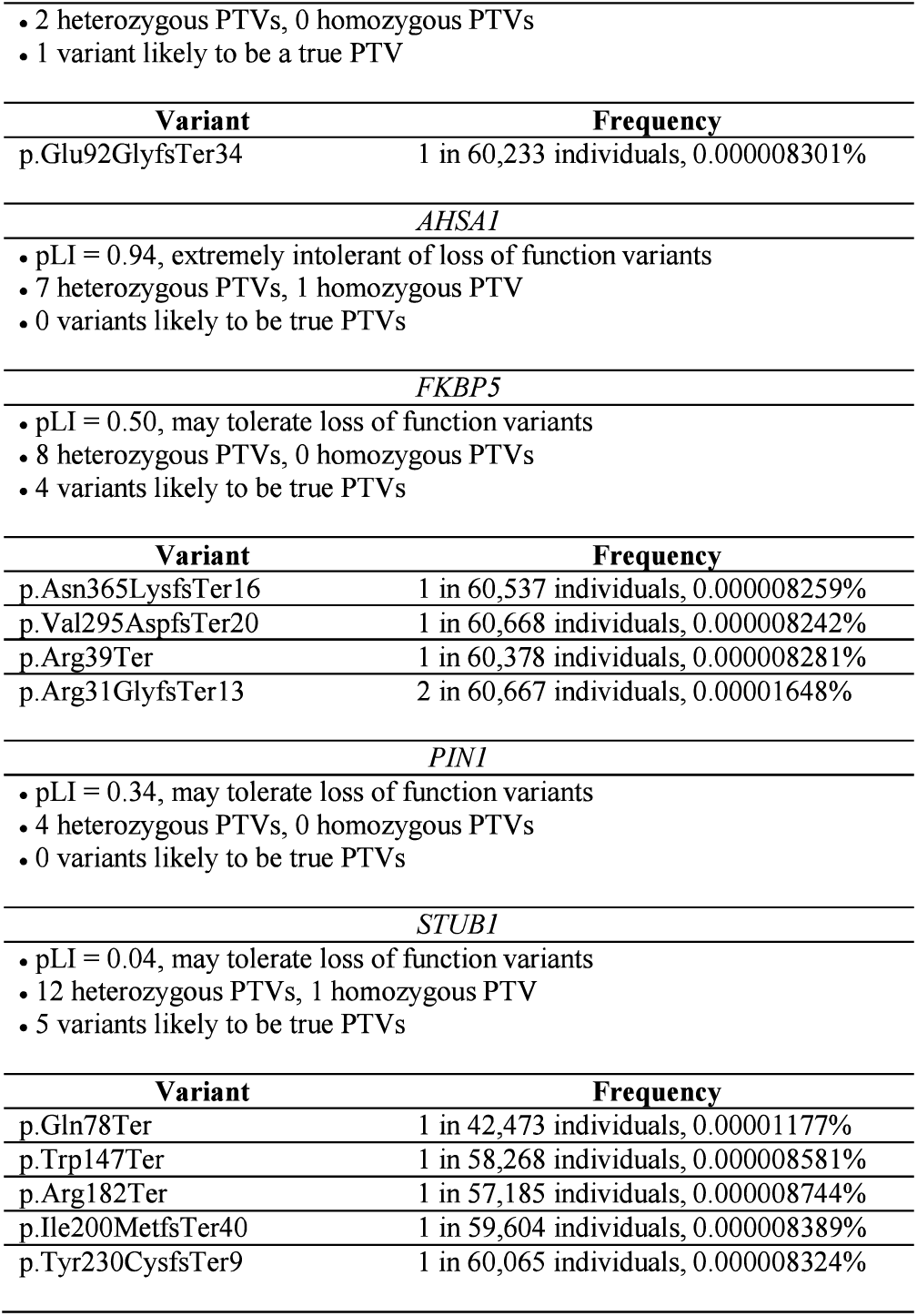
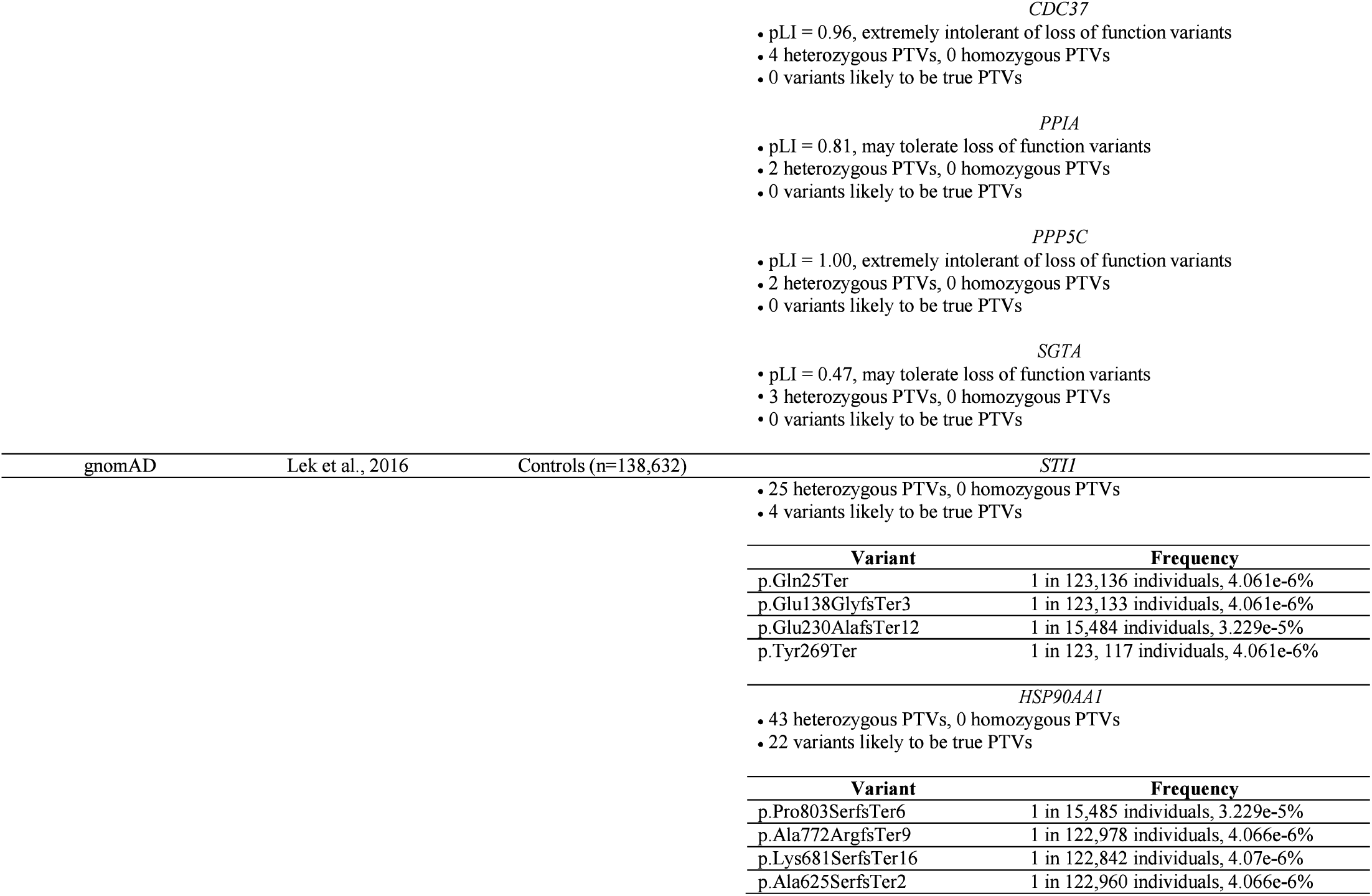
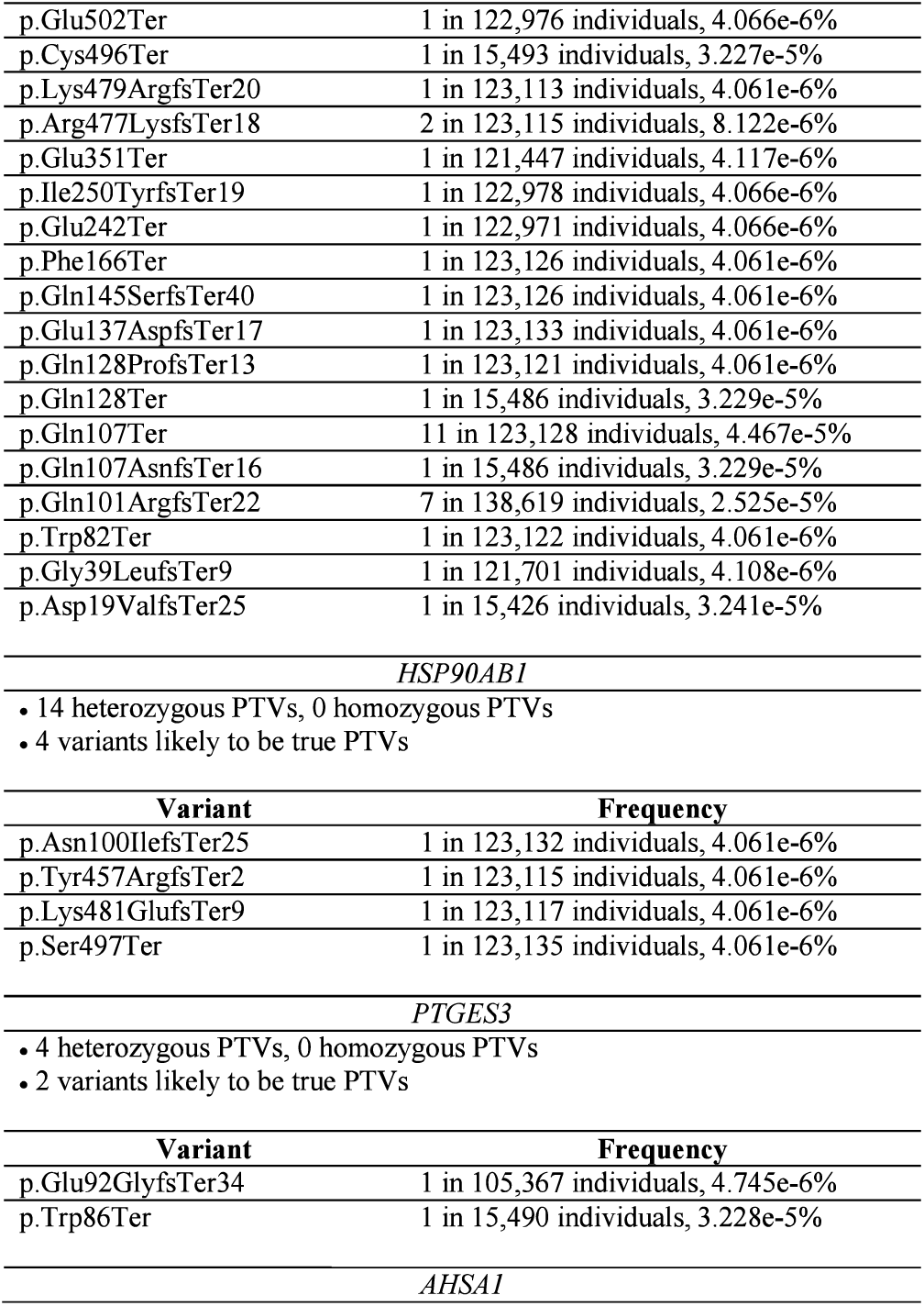
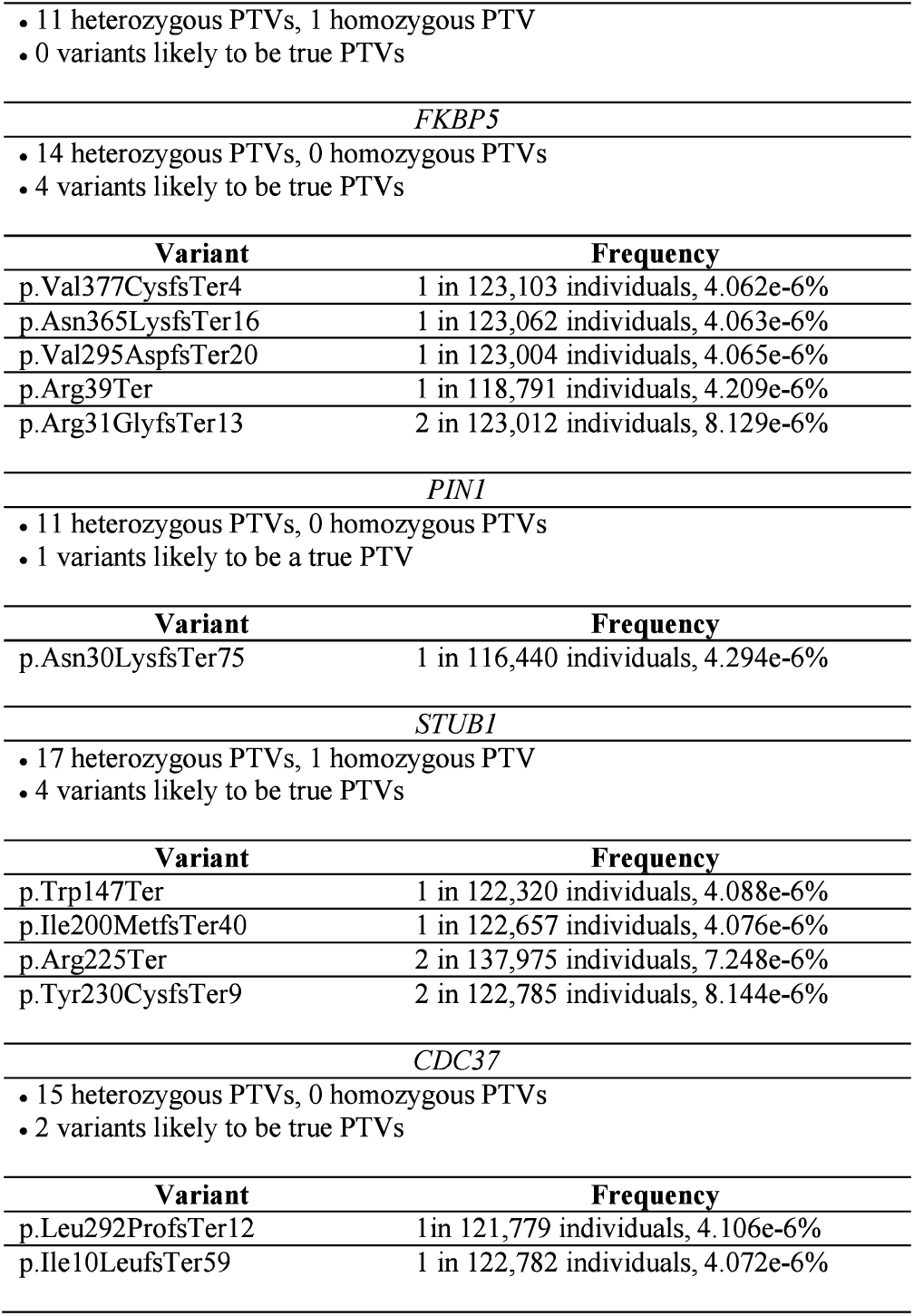
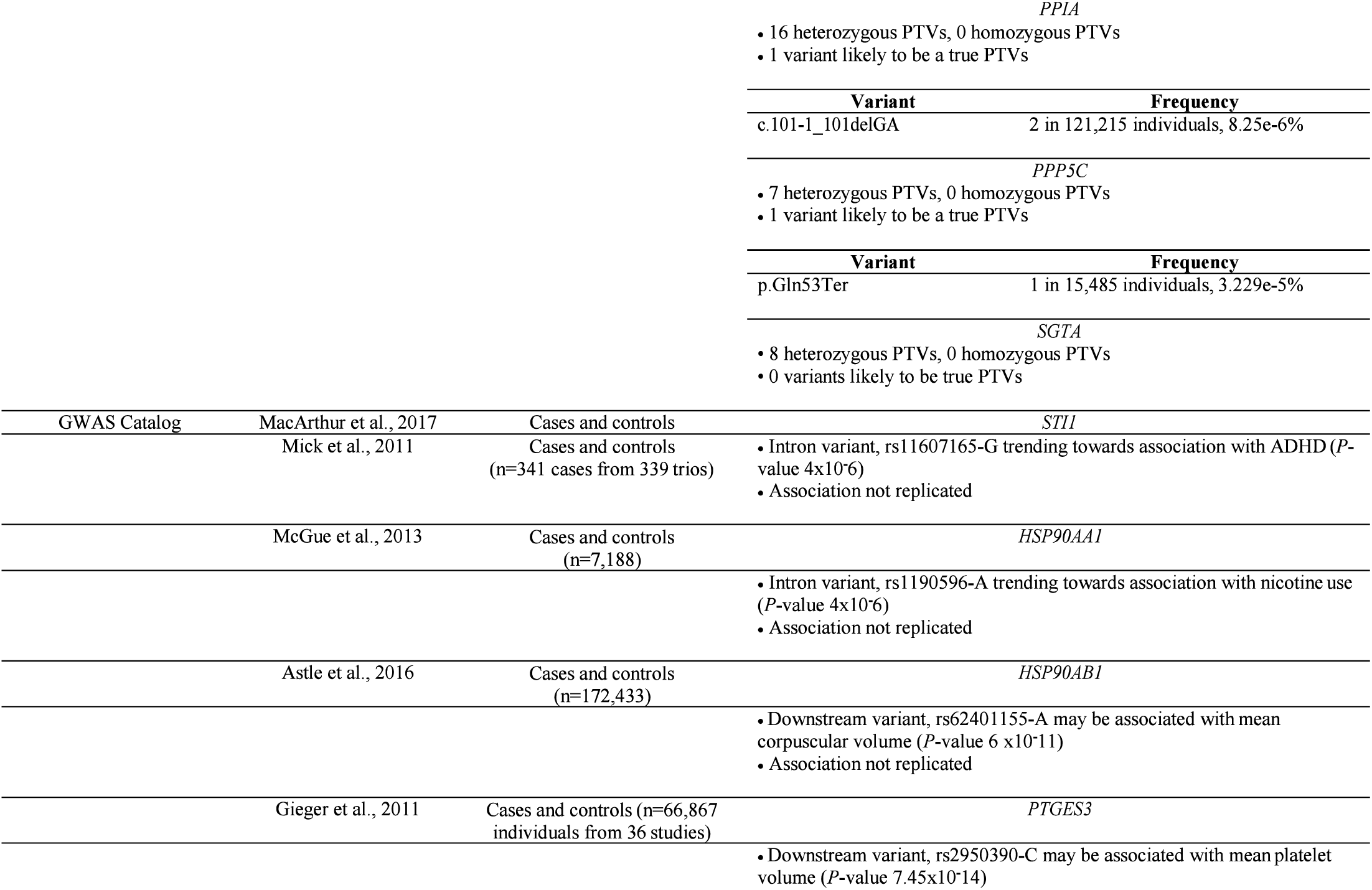
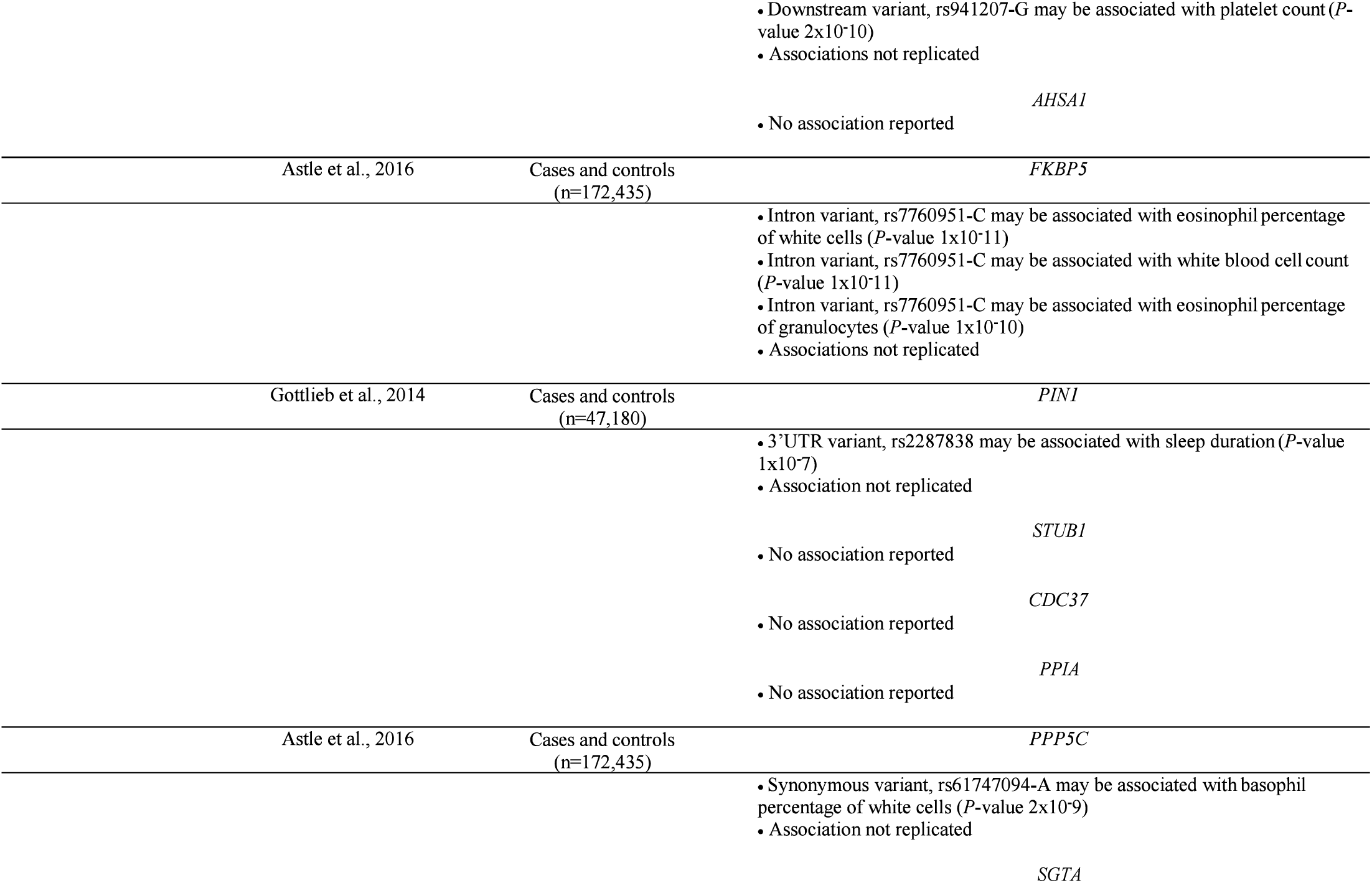
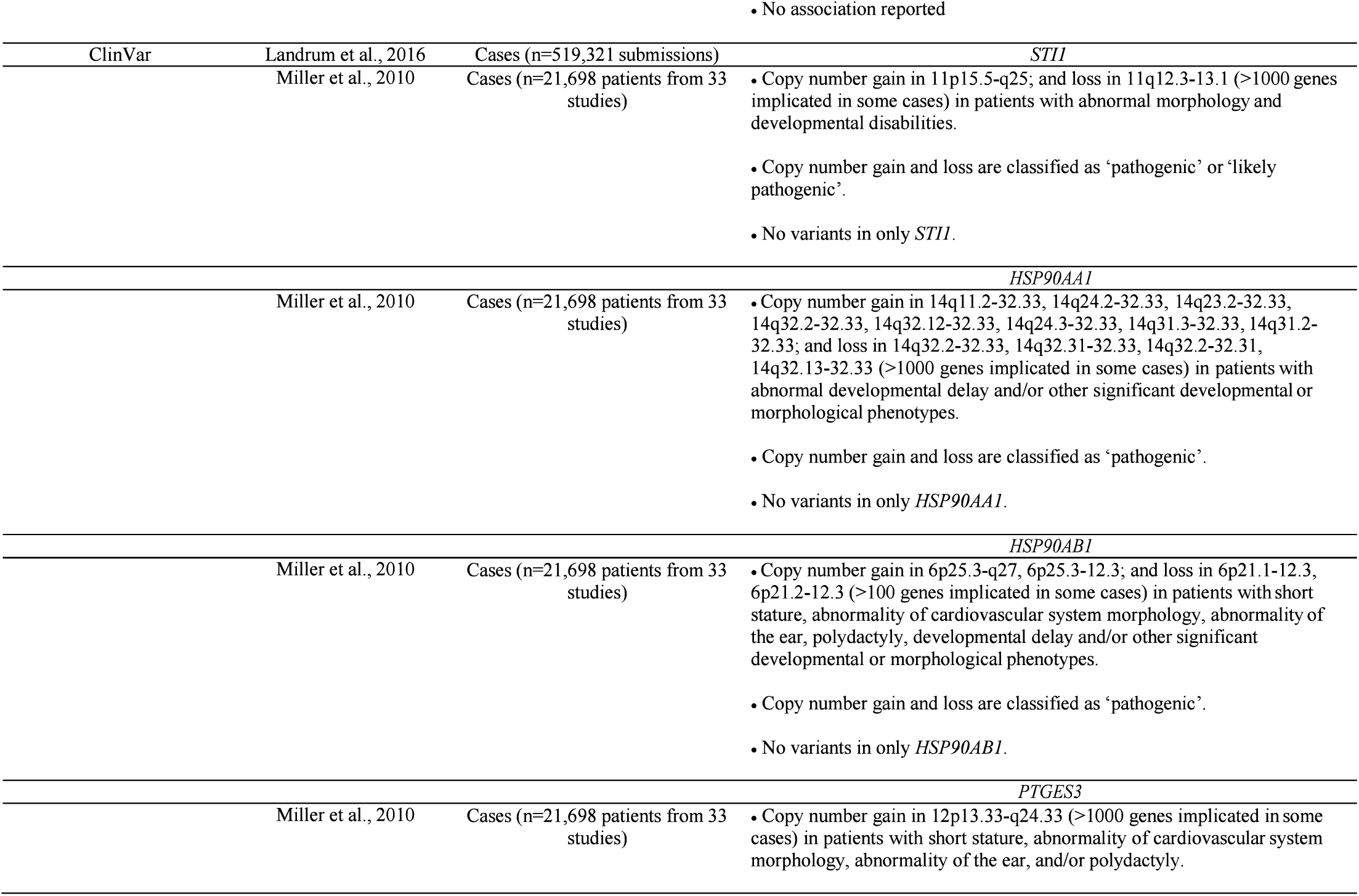
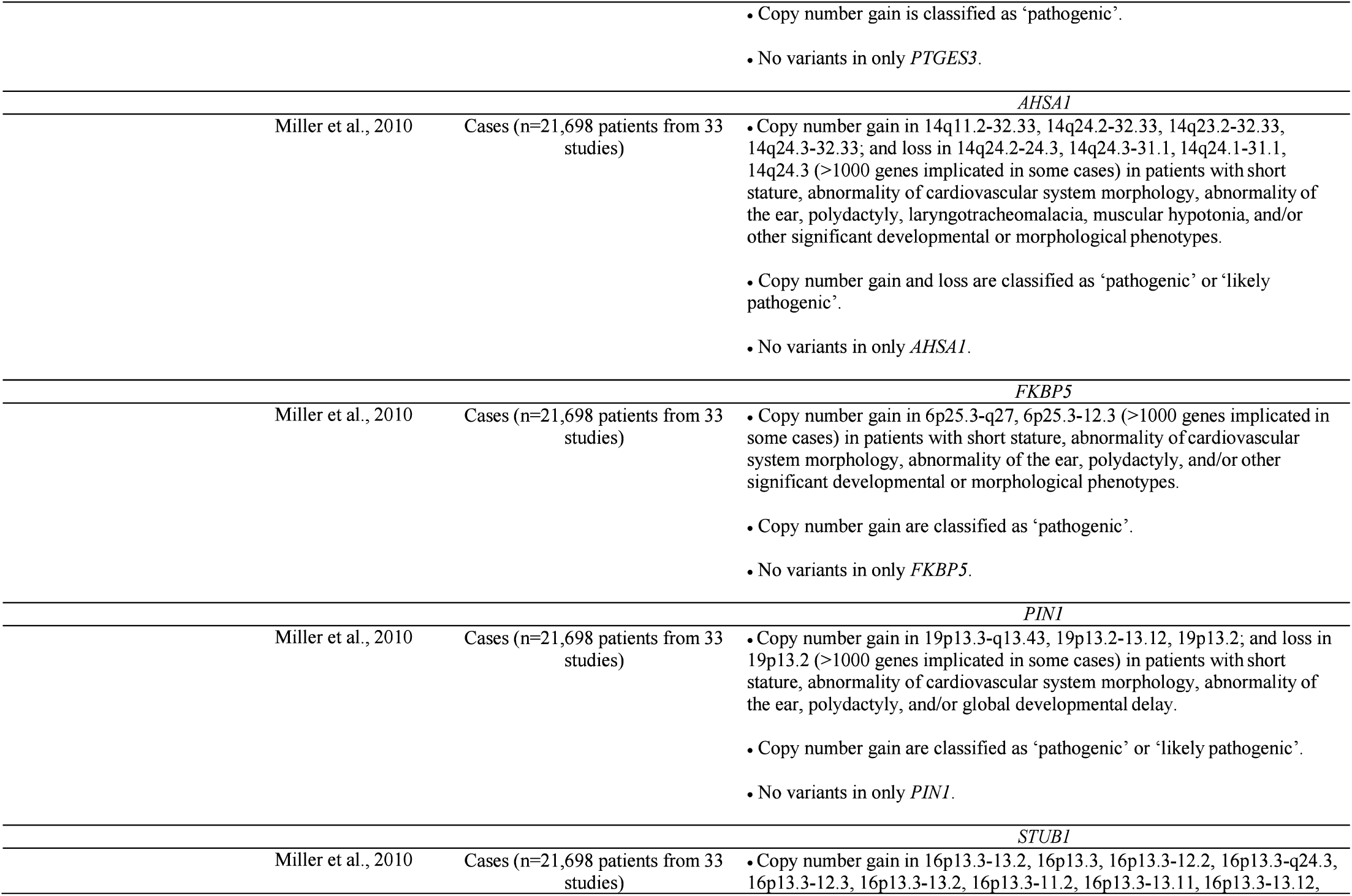
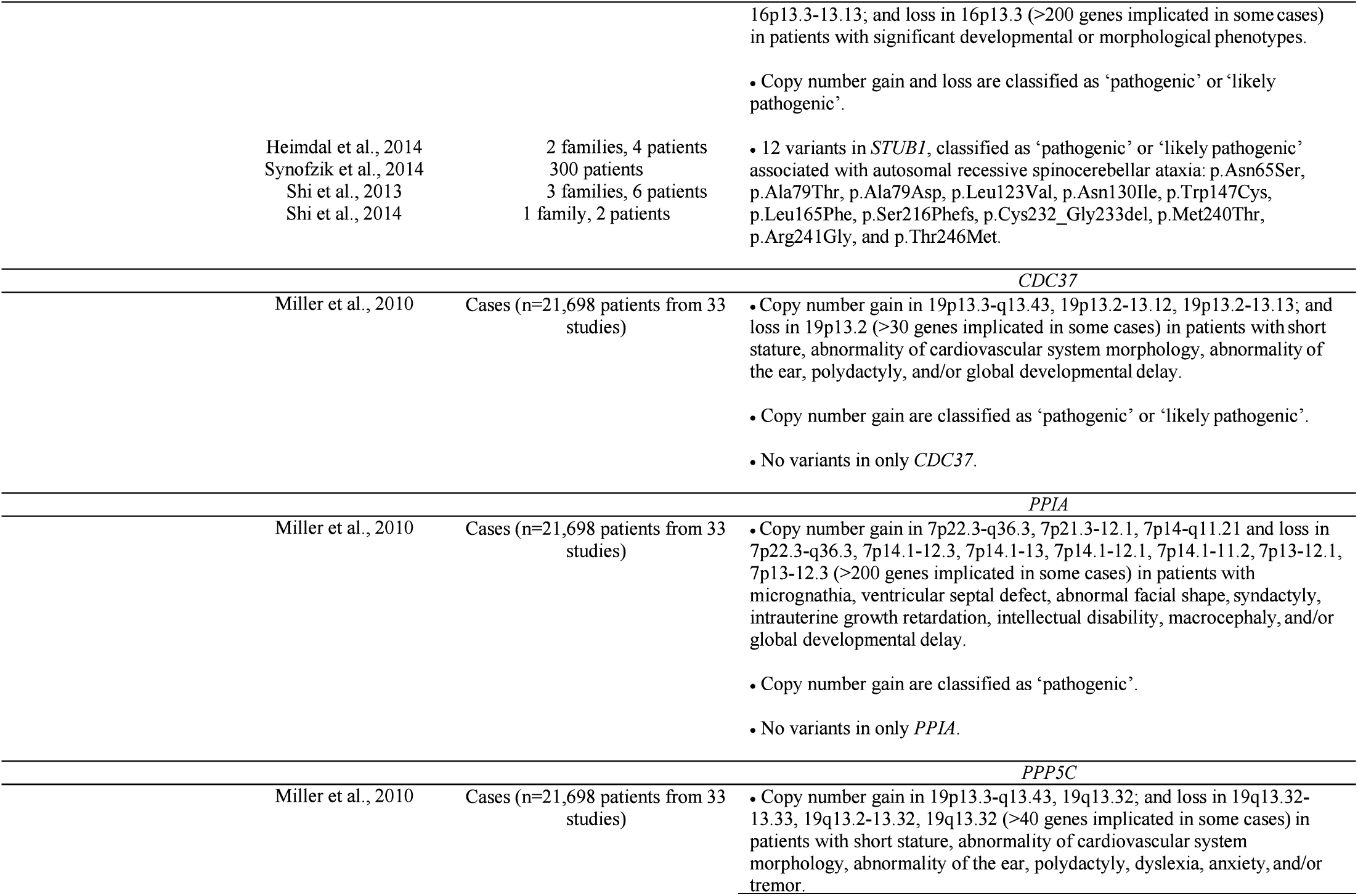
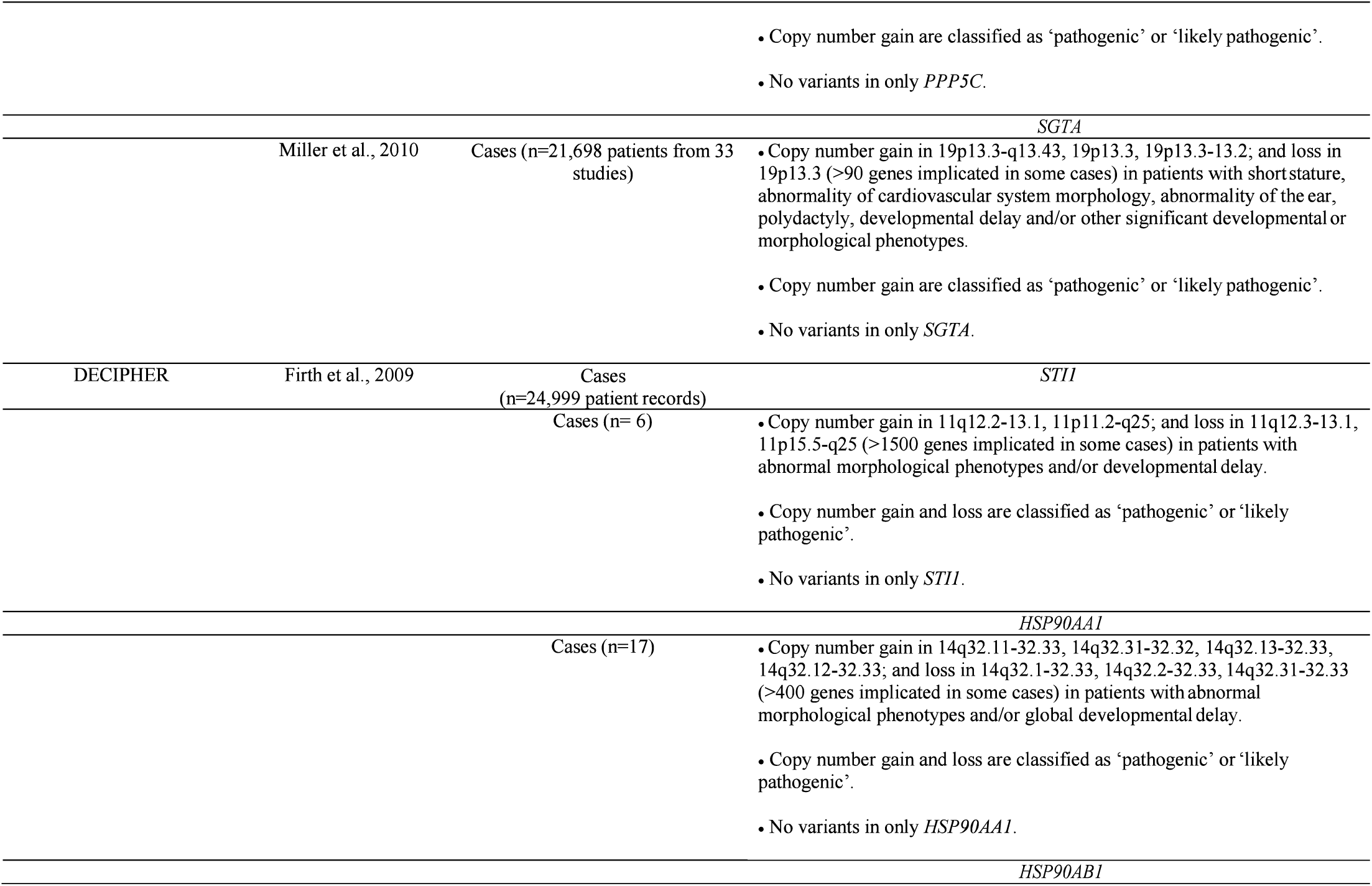
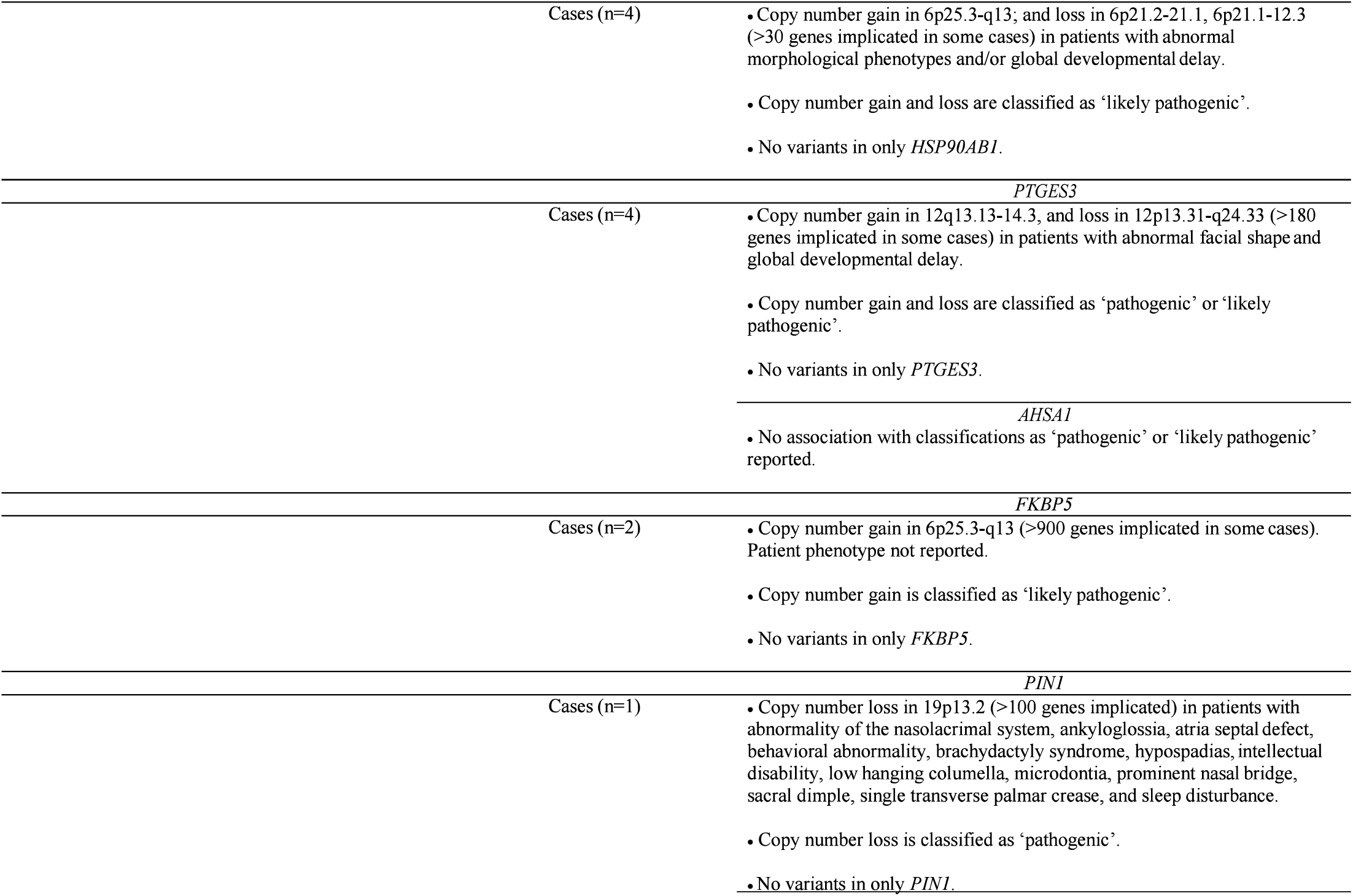
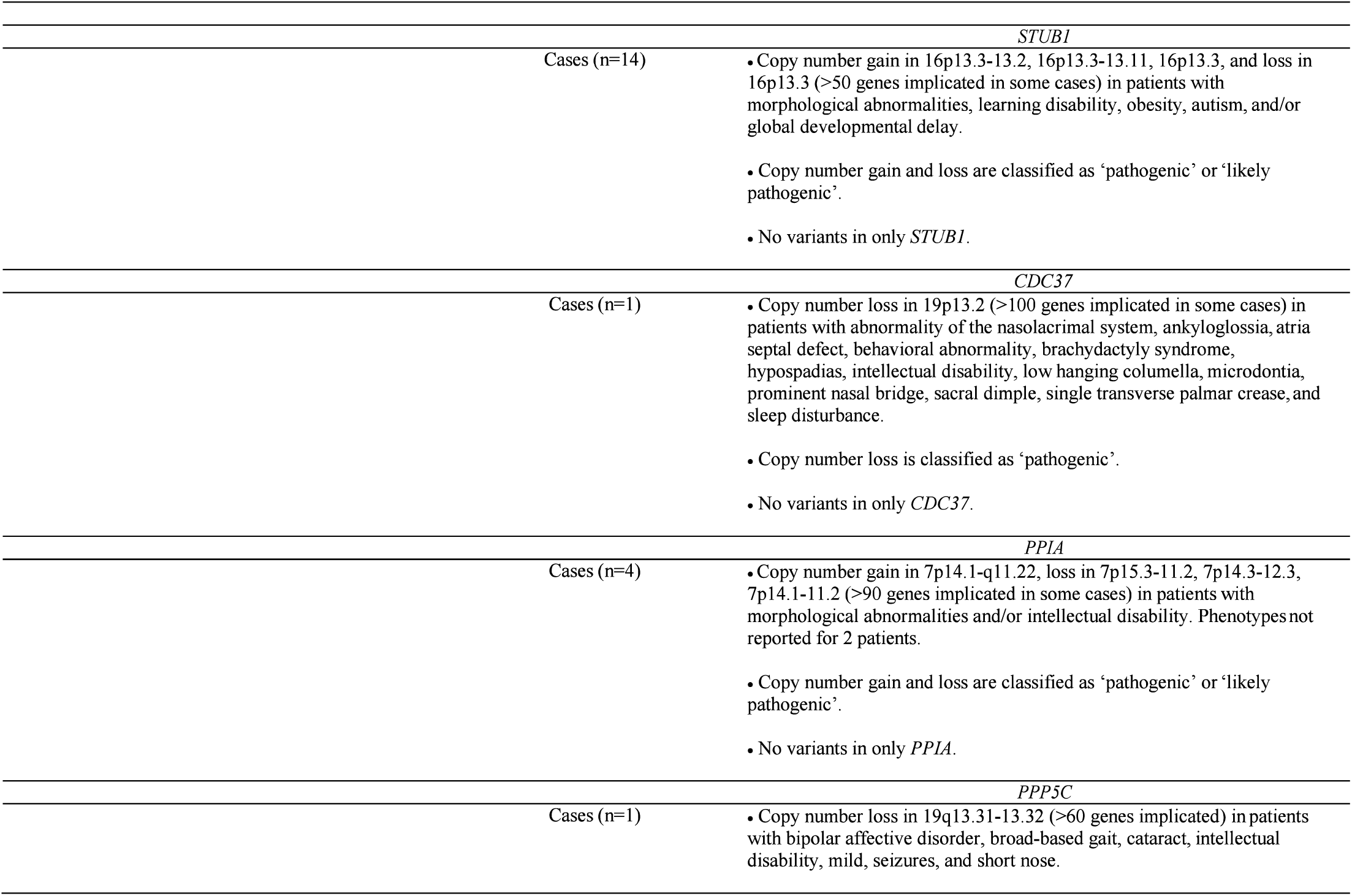
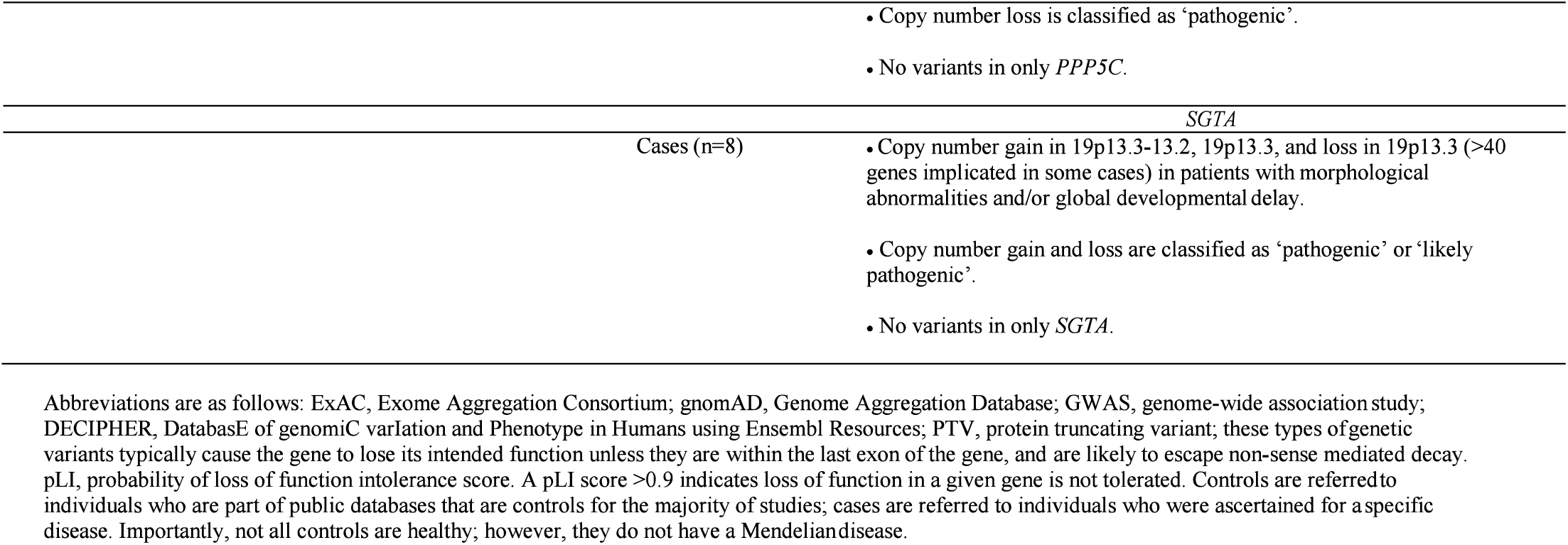
Analysis of probability of loss-of-function (pLI) score for *STIP1, HSP90AA1, HSP90AB1*, and other Hsp90 co-chaperones from publicly available databases for healthy and diseased patients. Genetic information of *STI1, HSP90AA1, HSP90AB1, PTGES3, AHSA1, FKBP5, PIN1, STUB1, CDC37, PPIA, PPP5C*, and *SGTA*, aggregated using publically available databases and repositories of healthy controls and disease-ascertained individuals.

**Supplementary Figure 1.**
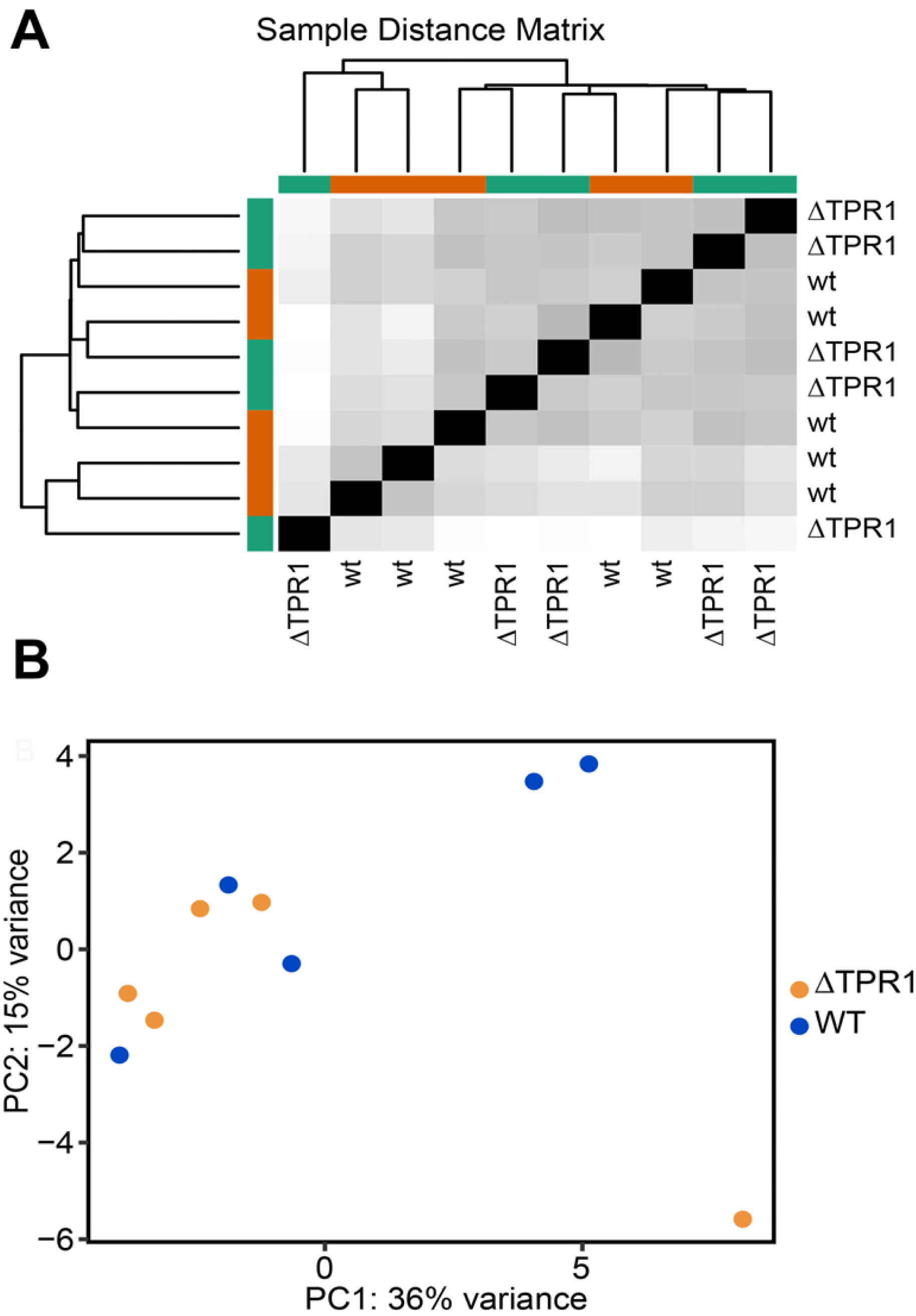
Unaltered Transcriptome in ΔTPR1 mice. Transcriptional changes between RNA samples from the cortex of 5 STI1 WT and 5 STI1 ΔTPR1 mice were analyzed by long RNA-sequencing. Bioinformatic analysis showed minimal variance between the samples in both **A)** sample distance matrix and **B)** principal component analysis.

## References

Adamowicz, D. H., Roy, S., Salmon, D. P., Galasko, D. R., Hansen, L. A., Masliah, E., & Gage, F. H. (2017). Hippocampal alpha-Synuclein in Dementia with Lewy Bodies Contributes to Memory Impairment and Is Consistent with Spread of Pathology. J Neurosci, 37(7), 1675–1684. doi:10.1523/JNEUROSCI.3047-16.2016

Afgan, E., Baker, D., van den Beek, M., Blankenberg, D., Bouvier, D., Cech, M., … Goecks, J. (2016). The Galaxy platform for accessible, reproducible and collaborative biomedical analyses: 2016 update. Nucleic Acids Res, 44(W1), W3–W10. doi:10.1093/nar/gkw343

Anders, S., Pyl, P. T., & Huber, W. (2015). HTSeq–a Python framework to work with high-throughput sequencing data. Bioinformatics, 31(2), 166–169. doi:10.1093/bioinformatics/btu638

Barent, R. L., Nair, S. C., Carr, D. C., Ruan, Y., Rimerman, R. A., Fulton, J., … Smith, D. F. (1998). Analysis of FKBP51/FKBP52 chimeras and mutants for Hsp90 binding and association with progesterone receptor complexes. Mol Endocrinol, 12(3), 342–354. doi:10.1210/mend.12.3.0075

Beers, M., & Kemphues, K. (2006). Depletion of the co-chaperone CDC-37 reveals two modes of PAR-6 cortical association in C. elegans embryos. Development, 133(19), 3745–3754. doi:10.1242/dev.02544

Beraldo, F. H., Ostapchenko, V. G., Caetano, F. A., Guimaraes, A. L., Ferretti, G. D., Daude, N., … Prado, M. A. (2016). Regulation of Amyloid beta Oligomer Binding to Neurons and Neurotoxicity by the Prion Protein-mGluR5 Complex. J Biol Chem, 291(42), 21945–21955. doi:10.1074/jbc.M116.738286

Beraldo, F. H., Soares, I. N., Goncalves, D. F., Fan, J., Thomas, A. A., Santos, T. G., … Prado, M. A. (2013). Stress-inducible phosphoprotein 1 has unique cochaperone activity during development and regulates cellular response to ischemia via the prion protein. FASEB J, 27(9), 3594–3607. doi:10.1096/fj.13-232280

Beraldo, F. H., Thomas, A., Kolisnyk, B., Hirata, P. H., De Jaeger, X., Martyn, A. C., … Prado, M. A. (2015). Hyperactivity and attention deficits in mice with decreased levels of stress-inducible phosphoprotein 1 (STIP1). Dis Model Mech, 8(11), 1457–1466. doi:10.1242/dmm.022525

Beyer, M. K., Bronnick, K. S., Hwang, K. S., Bergsland, N., Tysnes, O. B., Larsen, J. P., … Apostolova, L. G. (2013). Verbal memory is associated with structural hippocampal changes in newly diagnosed Parkinson’s disease. J Neurol Neurosurg Psychiatry, 84(1), 23–28. doi:10.1136/jnnp-2012-303054

Brehme, M., Voisine, C., Rolland, T., Wachi, S., Soper, J. H., Zhu, Y., … Morimoto, R. I. (2014). A chaperome subnetwork safeguards proteostasis in aging and neurodegenerative disease. Cell Rep, 9(3), 1135–1150. doi:10.1016/j.celrep.2014.09.042

Chang, H. C., Nathan, D. F., & Lindquist, S. (1997). In vivo analysis of the Hsp90 cochaperone Sti1 (p60). Mol Cell Biol, 17(1), 318–325.

Chen, L., Ding, Y., Cagniard, B., Van Laar, A. D., Mortimer, A., Chi, W., … Zhuang, X. (2008). Unregulated cytosolic dopamine causes neurodegeneration associated with oxidative stress in mice. J Neurosci, 28(2), 425–433. doi:10.1523/JNEUROSCI.3602-07.2008

Connarn, J. N., Assimon, V. A., Reed, R. A., Tse, E., Southworth, D. R., Zuiderweg, E. R., … Sun, D. (2014). The molecular chaperone Hsp70 activates protein phosphatase 5 (PP5) by binding the tetratricopeptide repeat (TPR) domain. J Biol Chem, 289(5), 2908–2917. doi:10.1074/jbc.M113.519421

Consortium, G. T. (2013). The Genotype-Tissue Expression (GTEx) project. Nat Genet, 45(6), 580–585. doi:10.1038/ng.2653

Cyr, D. M., Lu, X., & Douglas, M. G. (1992). Regulation of Hsp70 function by a eukaryotic DnaJ homolog. J Biol Chem, 267(29), 20927–20931.

Dai, Q., Zhang, C., Wu, Y., McDonough, H., Whaley, R. A., Godfrey, V., … Patterson, C. (2003). CHIP activates HSF1 and confers protection against apoptosis and cellular stress. Embo j, 22(20), 5446–5458. doi:10.1093/emboj/cdg529

Dickey, C. A., Kamal, A., Lundgren, K., Klosak, N., Bailey, R. M., Dunmore, J., … Petrucelli, L. (2007). The high-affinity HSP90-CHIP complex recognizes and selectively degrades phosphorylated tau client proteins. J Clin Invest, 117(3), 648–658. doi:10.1172/JCI29715

Ebong, I. O., Beilsten-Edmands, V., Patel, N. A., Morgner, N., & Robinson, C. V. (2016). The interchange of immunophilins leads to parallel pathways and different intermediates in the assembly of Hsp90 glucocorticoid receptor complexes. Cell Discov, 2, 16002. doi:10.1038/celldisc.2016.2

Ebrahimi-Fakhari, D., Saidi, L. J., & Wahlster, L. (2013). Molecular chaperones and protein folding as therapeutic targets in Parkinson’s disease and other synucleinopathies. Acta Neuropathol Commun, 1, 79. doi:10.1186/2051-5960-1- 79

Etoc, F., Metzger, J., Ruzo, A., Kirst, C., Yoney, A., Ozair, M. Z., … Siggia, E. D. (2016). A Balance between Secreted Inhibitors and Edge Sensing Controls Gastruloid Self-Organization. Dev Cell, 39(3), 302–315. doi:10.1016/j.devcel.2016.09.016

Farkas, Z., Kalapis, D., Bodi, Z., Szamecz, B., Daraba, A., Almasi, K., … Pal, C. (2018). Hsp70-associated chaperones have a critical role in buffering protein production costs. Elife, 7. doi:10.7554/eLife.29845

Farovik, A., Dupont, L. M., & Eichenbaum, H. (2010). Distinct roles for dorsal CA3 and CA1 in memory for sequential nonspatial events. Learn Mem, 17(1), 12–17. doi:10.1101/lm.1616209

Firth, H. V., Richards, S. M., Bevan, A. P., Clayton, S., Corpas, M., Rajan, D., … Carter, N. P. (2009). DECIPHER: Database of Chromosomal Imbalance and Phenotype in Humans Using Ensembl Resources. Am J Hum Genet, 84(4), 524–533. doi:10.1016/j.ajhg.2009.03.010

Floer, M., Bryant, G. O., & Ptashne, M. (2008). HSP90/70 chaperones are required for rapid nucleosome removal upon induction of the GAL genes of yeast. Proc Natl Acad Sci U S A, 105(8), 2975–2980. doi:10.1073/pnas.0800053105

Fontaine, S. N., Zheng, D., Sabbagh, J. J., Martin, M. D., Chaput, D., Darling, A., … Dickey, C. A. (2016). DnaJ/Hsc70 chaperone complexes control the extracellular release of neurodegenerative-associated proteins. Embo j, 35(14), 1537–1549. doi:10.15252/embj.201593489

Frydman, J., Nimmesgern, E., Ohtsuka, K., & Hartl, F. U. (1994). Folding of nascent polypeptide chains in a high molecular mass assembly with molecular chaperones. Nature, 370(6485), 111–117. doi:10.1038/370111a0

Fuhrmann-Stroissnigg, H., Ling, Y. Y., Zhao, J., McGowan, S. J., Zhu, Y., Brooks, R. W., … Robbins, P. D. (2017). Identification of HSP90 inhibitors as a novel class of senolytics. Nat Commun, 8(1), 422. doi:10.1038/s41467-017-00314-z

Gaiser, A. M., Brandt, F., & Richter, K. (2009a). The non-canonical Hop protein from Caenorhabditis elegans exerts essential functions and forms binary complexes with either Hsc70 or Hsp90. J Mol Biol, 391(3), 621–634. doi:10.1016/j.jmb.2009.06.051

Gaiser, A. M., Brandt, F., & Richter, K. (2009b). The Non-canonical Hop Protein from Caenorhabditis elegans Exerts Essential Functions and Forms Binary Complexes with Either Hsc70 or Hsp90. J. Mol. Biol.

Gangaraju, V. K., Yin, H., Weiner, M. M., Wang, J., Huang, X. A., & Lin, H. (2011). Drosophila Piwi functions in Hsp90-mediated suppression of phenotypic variation. Nat. Genet., 43(2), 153–158.

Gemmell, E., Bosomworth, H., Allan, L., Hall, R., Khundakar, A., Oakley, A. E., … Kalaria, R. N. (2012). Hippocampal neuronal atrophy and cognitive function in delayed poststroke and aging-related dementias. Stroke, 43(3), 808–814. doi:10.1161/STROKEAHA.111.636498

Gilbert, P. E., & Brushfield, A. M. (2009). The role of the CA3 hippocampal subregion in spatial memory: a process oriented behavioral assessment. Prog Neuropsychopharmacol Biol Psychiatry, 33(5), 774–781. doi:10.1016/j.pnpbp.2009.03.037

Grad, I., Cederroth, C. R., Walicki, J., Grey, C., Barluenga, S., Winssinger, N., … Picard, D. (2010). The molecular chaperone Hsp90alpha is required for meiotic progression of spermatocytes beyond pachytene in the mouse. PLoS One, 5(12), e15770.

Grad, I., McKee, T. A., Ludwig, S. M., Hoyle, G. W., Ruiz, P., Wurst, W., … Picard, D. (2006). The Hsp90 cochaperone p23 is essential for perinatal survival. Mol. Cell Biol., 26(23), 8976–8983.

Guzman, M. S., De Jaeger, X., Drangova, M., Prado, M. A., Gros, R., & Prado, V. F. (2013). Mice with selective elimination of striatal acetylcholine release are lean, show altered energy homeostasis and changed sleep/wake cycle. J Neurochem, 124(5), 658–669. doi:10.1111/jnc.12128

Guzman, M. S., De Jaeger, X., Raulic, S., Souza, I. A., Li, A. X., Schmid, S., … Prado, M. A. (2011). Elimination of the vesicular acetylcholine transporter in the striatum reveals regulation of behaviour by cholinergic-glutamatergic co-transmission. PLoS Biol, 9(11), e1001194. doi:10.1371/journal.pbio.1001194

Harst, A., Lin, H., & Obermann, W. M. (2005). Aha1 competes with Hop, p50 and p23 for binding to the molecular chaperone Hsp90 and contributes to kinase and hormone receptor activation. Biochem J, 387(Pt 3), 789–796. doi:10.1042/BJ20041283

Hildenbrand, Z. L., Molugu, S. K., Herrera, N., Ramirez, C., Xiao, C., & Bernal, R. A. (2011). Hsp90 can accommodate the simultaneous binding of the FKBP52 and HOP proteins. Oncotarget, 2(1-2), 43–58. doi:10.18632/oncotarget.225

Hoseini, H., Pandey, S., Jores, T., Schmitt, A., Franz-Wachtel, M., Macek, B., … Rapaport, D. (2016). The cytosolic cochaperone Sti1 is relevant for mitochondrial biogenesis and morphology. FEBS J, 283(18), 3338–3352. doi:10.1111/febs.13813

Janickova, H., Rosborough, K., Al-Onaizi, M., Kljakic, O., Guzman, M. S., Gros, R., … Prado, V. F. (2017). Deletion of the vesicular acetylcholine transporter from pedunculopontine/laterodorsal tegmental neurons modifies gait. J Neurochem, 140(5), 787–798. doi:10.1111/jnc.13910

Jinwal, U. K., Koren, J., 3rd, Borysov, S. I., Schmid, A. B., Abisambra, J. F., Blair, L. J., … Dickey, C. A. (2010). The Hsp90 cochaperone, FKBP51, increases Tau stability and polymerizes microtubules. J Neurosci, 30(2), 591–599. doi:10.1523/JNEUROSCI.4815-09.2010

Johnson, B. D., Schumacher, R. J., Ross, E. D., & Toft, D. O. (1998). Hop modulates Hsp70/Hsp90 interactions in protein folding. J Biol Chem, 273(6), 3679–3686.

Kalaitzakis, M. E., Christian, L. M., Moran, L. B., Graeber, M. B., Pearce, R. K., & Gentleman, S. M. (2009). Dementia and visual hallucinations associated with limbic pathology in Parkinson’s disease. Parkinsonism Relat Disord, 15(3), 196–204. doi:10.1016/j.parkreldis.2008.05.007

Karam, J. A., Parikh, R. Y., Nayak, D., Rosenkranz, D., & Gangaraju, V. K. (2017). Co-chaperone Hsp70/Hsp90-organizing protein (Hop) is required for transposon silencing and Piwi-interacting RNA (piRNA) biogenesis. J Biol Chem, 292(15), 6039–6046. doi:10.1074/jbc.C117.777730

Kim, D., Pertea, G., Trapnell, C., Pimentel, H., Kelley, R., & Salzberg, S. L. (2013). TopHat2: accurate alignment of transcriptomes in the presence of insertions, deletions and gene fusions. Genome Biol, 14(4), R36. doi:10.1186/gb-2013-14-4-r36

Kolisnyk, B., Al-Onaizi, M., Soreq, L., Barbash, S., Bekenstein, U., Haberman, N., … Prado, M. A. (2016). Cholinergic Surveillance over Hippocampal RNA Metabolism and Alzheimer’s-Like Pathology. Cereb Cortex. doi:10.1093/cercor/bhw177

Kolisnyk, B., Guzman, M. S., Raulic, S., Fan, J., Magalhaes, A. C., Feng, G., … Prado, M. A. (2013). ChAT-ChR2-EYFP mice have enhanced motor endurance but show deficits in attention and several additional cognitive domains. J Neurosci, 33(25), 10427–10438. doi:10.1523/JNEUROSCI.0395-13.2013

Kozak, M. (1986). Point mutations define a sequence flanking the AUG initiator codon that modulates translation by eukaryotic ribosomes. Cell, 44(2), 283–292.

Kuhn, H. G., Dickinson-Anson, H., & Gage, F. H. (1996). Neurogenesis in the dentate gyrus of the adult rat: age-related decrease of neuronal progenitor proliferation. J Neurosci, 16(6), 2027–2033.

Lackie, R. E., Maciejewski, A., Ostapchenko, V. G., Marques-Lopes, J., Choy, W. Y., Duennwald, M. L., … Prado, M. A. M. (2017). The Hsp70/Hsp90 Chaperone Machinery in Neurodegenerative Diseases. Front Neurosci, 11, 254. doi:10.3389/fnins.2017.00254

Landrum, M. J., Lee, J. M., Benson, M., Brown, G., Chao, C., Chitipiralla, S., … Maglott, D. R. (2016). ClinVar: public archive of interpretations of clinically relevant variants. Nucleic Acids Res, 44(D1), D862–868. doi:10.1093/nar/gkv1222

Lee, C. T., Graf, C., Mayer, F. J., Richter, S. M., & Mayer, M. P. (2012). Dynamics of the regulation of Hsp90 by the co-chaperone Sti1. Embo j, 31(6), 1518–1528. doi:10.1038/emboj.2012.37

Lee, I., Jerman, T. S., & Kesner, R. P. (2005). Disruption of delayed memory for a sequence of spatial locations following CA1- or CA3-lesions of the dorsal hippocampus. Neurobiol Learn Mem, 84(2), 138–147. doi:10.1016/j.nlm.2005.06.002

Lek, M., Karczewski, K. J., Minikel, E. V., Samocha, K. E., Banks, E., Fennell, T., … Exome Aggregation, C. (2016). Analysis of protein-coding genetic variation in 60,706 humans. Nature, 536(7616), 285–291. doi:10.1038/nature19057

Li, J., Richter, K., & Buchner, J. (2011). Mixed Hsp90-cochaperone complexes are important for the progression of the reaction cycle. Nat. Struct. Mol. Biol., 18(1), 61–66.

Li, J., Soroka, J., & Buchner, J. (2012). The Hsp90 chaperone machinery: Conformational dynamics and regulation by co-chaperones. Biochim. Biophys. Acta, 1823(3), 624–635.

Li, J. S., & Chao, Y. S. (2008). Electrolytic lesions of dorsal CA3 impair episodic-like memory in rats. Neurobiol Learn Mem, 89(2), 192–198. doi:10.1016/j.nlm.2007.06.006

Linden, R., Martins, V. R., Prado, M. A., Cammarota, M., Izquierdo, I., & Brentani, R. R. (2008). Physiology of the prion protein. Physiol Rev, 88(2), 673–728. doi:10.1152/physrev.00007.2007

Lopes, M. H., Hajj, G. N., Muras, A. G., Mancini, G. L., Castro, R. M., Ribeiro, K. C., … Martins, V. R. (2005). Interaction of cellular prion and stress-inducible protein 1 promotes neuritogenesis and neuroprotection by distinct signaling pathways. J Neurosci, 25(49), 11330–11339. doi:10.1523/JNEUROSCI.2313-05.2005

Love, M. I., Huber, W., & Anders, S. (2014). Moderated estimation of fold change and dispersion for RNA-seq data with DESeq2. Genome Biol, 15(12), 550. doi:10.1186/s13059-014-0550-8

MacArthur, J., Bowler, E., Cerezo, M., Gil, L., Hall, P., Hastings, E., … Parkinson, H. (2017). The new NHGRI-EBI Catalog of published genome-wide association studies (GWAS Catalog). Nucleic Acids Res, 45(D1), D896–D901. doi:10.1093/nar/gkw1133

Maciejewski, A., Ostapchenko, V. G., Beraldo, F. H., Prado, V. F., Prado, M. A., & Choy, W. Y. (2016). Domains of STIP 1 responsible for regulating PrPC-dependent amyloid-beta oligomer toxicity. Biochem J, 473(14), 2119–2130. doi:10.1042/BCJ20160087

MacLean, M., & Picard, D. (2003). Cdc37 goes beyond Hsp90 and kinases. Cell Stress. Chaperones., 8(2), 114–119.

Mayer, M. P. (2013). Hsp70 chaperone dynamics and molecular mechanism. Trends Biochem Sci, 38(10), 507–514. doi:10.1016/j.tibs.2013.08.001

Mick, E., McGough, J., Loo, S., Doyle, A. E., Wozniak, J., Wilens, T. E., … Faraone, S. V. (2011). Genome-wide association study of the child behavior checklist dysregulation profile. J Am Acad Child Adolesc Psychiatry, 50(8), 807–817 e808. doi:10.1016/j.jaac.2011.05.001

Migliorini, D., Lazzerini Denchi, E., Danovi, D., Jochemsen, A., Capillo, M., Gobbi, A., … Marine, J. C. (2002). Mdm4 (Mdmx) regulates p53-induced growth arrest and neuronal cell death during early embryonic mouse development. Mol Cell Biol, 22(15), 5527–5538.

Miller, D. T., Adam, M. P., Aradhya, S., Biesecker, L. G., Brothman, A. R., Carter, N. P., … Ledbetter, D. H. (2010). Consensus statement: chromosomal microarray is a first-tier clinical diagnostic test for individuals with developmental disabilities or congenital anomalies. Am J Hum Genet, 86(5), 749–764. doi:10.1016/j.ajhg.2010.04.006

Morgner, N., Schmidt, C., Beilsten-Edmands, V., Ebong, I. O., Patel, N. A., Clerico, E. M., … Robinson, C. V. (2015). Hsp70 forms antiparallel dimers stabilized by post-translational modifications to position clients for transfer to Hsp90. Cell Rep, 11(5), 759–769. doi:10.1016/j.celrep.2015.03.063

Nair, S. C., Rimerman, R. A., Toran, E. J., Chen, S., Prapapanich, V., Butts, R. N., & Smith, D. F. (1997). Molecular cloning of human FKBP51 and comparisons of immunophilin interactions with Hsp90 and progesterone receptor. Mol Cell Biol, 17(2), 594–603.

Nicolet, C. M., & Craig, E. A. (1989). Isolation and characterization of STI1, a stress-inducible gene from Saccharomyces cerevisiae. Mol Cell Biol, 9(9), 3638–3646.

Ostapchenko, V. G., Beraldo, F. H., Mohammad, A. H., Xie, Y. F., Hirata, P. H., Magalhaes, A. C., … Prado, M. A. (2013). The prion protein ligand, stress-inducible phosphoprotein 1, regulates amyloid-beta oligomer toxicity. J Neurosci, 33(42), 16552–16564. doi:10.1523/JNEUROSCI.3214-13.2013

Padurariu, M., Ciobica, A., Mavroudis, I., Fotiou, D., & Baloyannis, S. (2012). Hippocampal neuronal loss in the CA1 and CA3 areas of Alzheimer’s disease patients. Psychiatr Danub, 24(2), 152–158.

Philp, L. K., Day, T. K., Butler, M. S., Laven-Law, G., Jindal, S., Hickey, T. E., … Tilley, W. D. (2016). Small Glutamine-Rich Tetratricopeptide Repeat-Containing Protein Alpha (SGTA) Ablation Limits Offspring Viability and Growth in Mice. Sci Rep, 6, 28950. doi:10.1038/srep28950

Picard, D. (2006). Chaperoning steroid hormone action. Trends Endocrinol. Metab, 17(6), 229–235.

Pratt, W. B., Gestwicki, J. E., Osawa, Y., & Lieberman, A. P. (2015). Targeting Hsp90/Hsp70-based protein quality control for treatment of adult onset neurodegenerative diseases. Annu Rev Pharmacol Toxicol, 55, 353–371. doi:10.1146/annurev-pharmtox-010814-124332

Retzlaff, M., Hagn, F., Mitschke, L., Hessling, M., Gugel, F., Kessler, H., … Buchner, J. (2010). Asymmetric activation of the hsp90 dimer by its cochaperone aha1. Mol. Cell, 37(3), 344–354.

Robitsek, J., Ratner, M. H., Stewart, T., Eichenbaum, H., & Farb, D. H. (2015). Combined administration of levetiracetam and valproic acid attenuates age-related hyperactivity of CA3 place cells, reduces place field area, and increases spatial information content in aged rat hippocampus. Hippocampus, 25(12), 1541–1555. doi:10.1002/hipo.22474

Rodina, A., Wang, T., Yan, P., Gomes, E. D., Dunphy, M. P., Pillarsetty, N., … Chiosis, G. (2016). The epichaperome is an integrated chaperome network that facilitates tumour survival. Nature, 538(7625), 397–401. doi:10.1038/nature19807

Rohl, A., Tippel, F., Bender, E., Schmid, A. B., Richter, K., Madl, T., & Buchner, J. (2015). Hop/Sti1 phosphorylation inhibits its co-chaperone function. EMBO Rep, 16(2), 240–249. doi:10.15252/embr.201439198

Rohl, A., Wengler, D., Madl, T., Lagleder, S., Tippel, F., Herrmann, M., … Buchner, J. (2015). Hsp90 regulates the dynamics of its cochaperone Sti1 and the transfer of Hsp70 between modules. Nat Commun, 6, 6655. doi:10.1038/ncomms7655

Roy, A., Fields, W. C., Rocha-Resende, C., Resende, R. R., Guatimosim, S., Prado, V. F., … Prado, M. A. (2013). Cardiomyocyte-secreted acetylcholine is required for maintenance of homeostasis in the heart. FASEB J, 27(12), 5072–5082. doi:10.1096/fj.13-238279

Sahasrabudhe, P., Rohrberg, J., Biebl, M. M., Rutz, D. A., & Buchner, J. (2017). The Plasticity of the Hsp90 Co-chaperone System. Mol Cell, 67(6), 947–961 e945. doi:10.1016/j.molcel.2017.08.004

Santpere, G., Garcia-Esparcia, P., Andres-Benito, P., Lorente-Galdos, B., Navarro, A., & Ferrer, I. (2018). Transcriptional network analysis in frontal cortex in Lewy body diseases with focus on dementia with Lewy bodies. Brain Pathol, 28(3), 315–333. doi:10.1111/bpa.12511

Sawarkar, R., Sievers, C., & Paro, R. (2012). Hsp90 globally targets paused RNA polymerase to regulate gene expression in response to environmental stimuli. Cell, 149(4), 807–818.

Schmid, A. B., Lagleder, S., Grawert, M. A., Rohl, A., Hagn, F., Wandinger, S. K., … Buchner, J. (2012a). The architecture of functional modules in the Hsp90 co-chaperone Sti1/Hop. EMBO J.

Schmid, A. B., Lagleder, S., Grawert, M. A., Rohl, A., Hagn, F., Wandinger, S. K., … Buchner, J. (2012b). The architecture of functional modules in the Hsp90 co-chaperone Sti1/Hop. Embo j, 31(6), 1506–1517. doi:10.1038/emboj.2011.472

Soares, I. N., Caetano, F. A., Pinder, J., Rodrigues, B. R., Beraldo, F. H., Ostapchenko, V. G., … Prado, M. A. (2013). Regulation of stress-inducible phosphoprotein 1 nuclear retention by protein inhibitor of activated STAT PIAS1. Mol Cell Proteomics, 12(11), 3253–3270. doi:10.1074/mcp.M113.031005

Song, H. O., Lee, W., An, K., Lee, H. S., Cho, J. H., Park, Z. Y., & Ahnn, J. (2009). C. elegans STI-1, the homolog of Sti1/Hop, is involved in aging and stress response. J Mol Biol, 390(4), 604–617. doi:10.1016/j.jmb.2009.05.035

Song, Y., & Masison, D. C. (2005). Independent regulation of Hsp70 and Hsp90 chaperones by Hsp70/Hsp90-organizing protein Sti1 (Hop1). J Biol Chem, 280(40), 34178–34185. doi:10.1074/jbc.M505420200

Steffenach, H. A., Sloviter, R. S., Moser, E. I., & Moser, M. B. (2002). Impaired retention of spatial memory after transection of longitudinally oriented axons of hippocampal CA3 pyramidal cells. Proc Natl Acad Sci U S A, 99(5), 3194–3198. doi:10.1073/pnas.042700999

Stenson, P. D., Mort, M., Ball, E. V., Evans, K., Hayden, M., Heywood, S., … Cooper, D. N. (2017). The Human Gene Mutation Database: towards a comprehensive repository of inherited mutation data for medical research, genetic diagnosis and next-generation sequencing studies. Hum Genet, 136(6), 665–677. doi:10.1007/s00439-017-1779-6

Taipale, M., Krykbaeva, I., Koeva, M., Kayatekin, C., Westover, K. D., Karras, G. I., & Lindquist, S. (2012). Quantitative analysis of HSP90-client interactions reveals principles of substrate recognition. Cell, 150(5), 987–1001. doi:10.1016/j.cell.2012.06.047

Team, R. (2017). R: A language and environment for statistical computing. R Foundation for Statistical computing.

Tsai, J., & Douglas, M. G. (1996). A conserved HPD sequence of the J-domain is necessary for YDJ1 stimulation of Hsp70 ATPase activity at a site distinct from substrate binding. J Biol Chem, 271(16), 9347–9354.

Voss, A. K., Thomas, T., & Gruss, P. (2000). Mice lacking HSP90beta fail to develop a placental labyrinth. Development, 127(1), 1–11.

Wandinger, S. K., Suhre, M. H., Wegele, H., & Buchner, J. (2006). The phosphatase Ppt1 is a dedicated regulator of the molecular chaperone Hsp90. Embo j, 25(2), 367–376. doi:10.1038/sj.emboj.7600930

Wolfe, K. J., Ren, H. Y., Trepte, P., & Cyr, D. M. (2013). The Hsp70/90 cochaperone, Sti1, suppresses proteotoxicity by regulating spatial quality control of amyloid-like proteins. Mol Biol Cell, 24(23), 3588–3602. doi:10.1091/mbc.E13-06-0315

Yong, W., Bao, S., Chen, H., Li, D., Sanchez, E. R., & Shou, W. (2007). Mice lacking protein phosphatase 5 are defective in ataxia telangiectasia mutated (ATM)-mediated cell cycle arrest. J Biol Chem, 282(20), 14690–14694. doi:10.1074/jbc.C700019200

Yong, W., Yang, Z., Periyasamy, S., Chen, H., Yucel, S., Li, W., … Shou, W. (2007). Essential role for Co-chaperone Fkbp52 but not Fkbp51 in androgen receptor-mediated signaling and physiology. J Biol Chem, 282(7), 5026–5036. doi:10.1074/jbc.M609360200

Zanata, S. M., Lopes, M. H., Mercadante, A. F., Hajj, G. N., Chiarini, L. B., Nomizo, R., … Martins, V. R. (2002). Stress - inducible protein 1 is a cell surface ligand for cellular prion that triggers neuroprotection. Embo j, 21(13), 3307–3316. doi:10.1093/emboj/cdf325

Zhang, M., Boter, M., Li, K., Kadota, Y., Panaretou, B., Prodromou, C., … Pearl, L. H. (2008). Structural and functional coupling of Hsp90- and Sgt1-centred multi-protein complexes. Embo j, 27(20), 2789–2798. doi:10.1038/emboj.2008.190

Zhao, R., Davey, M., Hsu, Y. C., Kaplanek, P., Tong, A., Parsons, A. B., … Houry, W. A. (2005). Navigating the chaperone network: an integrative map of physical and genetic interactions mediated by the hsp90 chaperone. Cell, 120(5), 715–727. doi:10.1016/j.cell.2004.12.024

Zhou, H., Huang, C., Chen, H., Wang, D., Landel, C. P., Xia, P. Y., … Xia, X. G. (2010). Transgenic rat model of neurodegeneration caused by mutation in the TDP gene. PLoS Genet, 6(3), e1000887. doi:10.1371/journal.pgen.1000887

